# Timing of transcriptomic peripheral blood mononuclear cell responses of sheep to *Fasciola hepatica* infection differs from those of cattle, reflecting different disease phenotypes

**DOI:** 10.1101/2021.06.21.449266

**Authors:** Dagmara A. Niedziela, Amalia Naranjo-Lucena, Verónica Molina-Hernández, John A. Browne, Álvaro Martínez-Moreno, José Pérez, David E. MacHugh, Grace Mulcahy

## Abstract

Infection with the zoonotic trematode *Fasciola hepatica*, common in many regions with a temperate climate, leads to delayed growth and loss of productivity in cattle, while infection in sheep can have more severe effects, potentially leading to death. Previous transcriptomic analyses revealed upregulation of *TGFB1*, cell death and Toll-like receptor signalling, T-cell activation, and inhibition of nitric oxide production in macrophages in response to infection. However, the differences between ovine and bovine responses have not yet been explored. The objective of this study was to further investigate the transcriptomic response of ovine peripheral blood mononuclear cells (PBMC) to *F. hepatica* infection, and to elucidate the differences between ovine and bovine PBMC responses.

Sixteen male Merino sheep were randomly assigned to infected or control groups (n = 8 per group) and orally infected with 120 *F. hepatica* metacercariae. Transcriptomic data was generated from PBMC at 0, 2 and 16 weeks post-infection (wpi), and analysed for differentially expressed (DE) genes between infected and control animals at each time point (analysis 1), and for each group relative to time 0 (analysis 2). Analysis 2 was then compared to a similar study performed previously on bovine PBMC.

A total of 453 DE genes were found at 2 wpi, and 2 DE genes at 16 wpi (FDR < 0.1, analysis 1). Significantly overrepresented biological pathways at 2 wpi included *role of PKR in interferon induction and anti-viral response*, *death receptor signalling* and *RIG-I-like receptor signalling*, which suggested that an activation of innate response to intracellular nucleic acids and inhibition of cellular apoptosis were taking place. Comparison of analysis 2 with the previous bovine transcriptomic study revealed that anti-inflammatory response pathways which were significantly overrepresented in the acute phase in cattle, including *IL-10 signalling*, *Th2 pathway*, and *Th1 and Th2 activation* were upregulated only in the chronic phase in sheep. We propose that the earlier activation of anti-inflammatory responses in cattle, as compared with sheep, may be related to the general absence of acute clinical signs in cattle. These findings offer scope for “smart vaccination” strategies for this important livestock parasite.

## 1 Introduction

*Fasciola hepatica* is a zoonotic trematode that affects livestock (sheep, cattle, goats) as well as humans. Human cases of fasciolosis occur mainly in Latin America, Africa and Asia, with approximately 2.5 million people affected worldwide (1; 2). Sporadic human cases also occur in Europe as a result of the consumption of raw vegetables such as parsley or watercress, grown on irrigated or flooded land (3).

*F. hepatica* infection is an important animal health issue in ruminants in endemic areas (4). In Ireland, *F. hepatica* seroprevalence is reported as 75.4% to 82% of dairy herds, and within-herd prevalence is estimated at ≤□50% (5; 6). In Northern Ireland, 63.15% cattle herds were positive for *F. hepatica*, with an overall animal level prevalence of 23.68% (7). Horses can also become infected, with a 9.5% prevalence of *F. hepatica* antibodies reported in a post-mortem survey of horses in Ireland (8). The cost of livestock infection includes productivity losses (e.g., meat, milk, wool) as well as reduced fertility and condemnation of livers as unfit for human consumption (9; 10). While *F. hepatica* infection in cattle leads to delayed growth and loss of productivity, acute infection in sheep can have more severe effects. The severe disease presentation in sheep is linked to a rapid acute response and can lead to sudden death (11). Anthelmintic resistance in *F. hepatica* populations is an increasingly important issue, with resistance to triclabendazole of particular concern (12; 13), as this is the only drug effective against early immature flukes. Hence, there has been significant attention given to developing a vaccine effective in protecting ruminants against fasciolosis. While promising vaccine trials have been conducted, repeatability and consistency has been an issue (14; 15).

The ruminant immune response to *F. hepatica* infection involves an increase in peripheral eosinophil numbers, alternative activation of macrophages, and suppression of Th1 responsiveness to the parasites themselves and to bystander antigens (16; 17). The transcriptomic response of peripheral blood mononuclear cells (PBMC) to *F. hepatica* in cattle has been characterised by apoptosis of antigen-presenting cells (APCs), liver fibrosis and a Th2 response, with *TNF*, *IL1B*, and *DUSP1, APP, STAT3* and *mir-155* as important upstream regulator genes leading to hepatic fibrosis and apoptosis of APCs or migration and chemotaxis of leukocytes (18). A balanced Treg-Th2 response was present in the acute phase with increased polarization towards a Th2 response during the chronic phase of infection.

The transcriptomic response of ovine PBMC has also been characterised previously. The response at 2 and 8 weeks post-infection (wpi) was associated with upregulation of the TGF-β signalling pathway, the complement system, chemokine signalling and T-cell activation. Groups of genes were identified which were important for the immune response to the parasite and included those coding for lectins, pro-inflammatory molecules from the S100 family and transmembrane glycoproteins from the CD300 family. Early infection (2 wpi) was characterised by positive activation of T-cell migration and leukocyte activation, while the in the later stage (8 wpi) lipoxin metabolic processes and Fc gamma R-mediated phagocytosis were among the upregulated pathways (19). Another study which looked at the ovine acute and chronic response at 1 and 14 wpi identified upregulation of TGFβ signalling, production of nitric oxide in macrophages, death receptor signalling and IL-17 signalling at 2 wpi, as well as Toll-like receptor (TLR) and p53 signalling in both the acute and chronic phase (20). The overall conclusions of these studies suggest a drive towards an anti-inflammatory response and tissue repair in *F. hepatica* infection in both sheep and cattle.

The objective of this study was to further investigate the pathways involved in the response of ovine PBMC to *F. hepatica* infection, and to compare those with bovine responses to aid in developing effective vaccine strategies.

## 2 Materials and methods

### 2.1 Animal trial outline

The animal trial has been described previously by (21). Briefly, nineteen male Merino sheep were obtained from a liver fluke-free farm at 9 months of age. The animals were all purchased into the research farm at the same time and were acclimatized for 3 months before the infections. All animals were tested monthly for parasite eggs by faecal sedimentation. Prior to challenge, all animals were tested for serum IgG specific for FhCL1 by an enzyme-linked immunosorbent assay (ELISA). Animals were housed indoors (100 m^2^ covered and 100 m^2^ uncovered facilities) and fed with hay and pellets and given water *ad libitum.* The infected and control animals were allocated in the same house (co-housed). Sheep were randomly allocated to the infected and control group, with 11 animals in the infected group, and 8 animals in the control group. However, 3 infected animals were later excluded from the study due to: 1) sample not being available at 16 wpi (not enough PBMC), 2) an animal identified with a potential pre-existing condition due to high eosinophils pre-infection (9.2 %), low haemoglobin (anaemic), and liver atrophy on pathology exam, and 3) a potentially mislabelled sample. This resulted in *n* = 8 in both groups. At week 0, eight sheep were orally infected with one dose of 120 *F. hepatica* metacercariae of the Ridgeway (South Gloucester, UK) strain. Animals were monitored for 16 weeks, with faecal egg counts measured weekly starting from 7 wpi, and haematology measurements taken at 0, 2, 7 and 16 wpi. PMBC were purified from blood samples taken at 0, 2 and 16 wpi and used for the transcriptomic analysis (**Fig. 1A**). Sheep were sacrificed at 16 wpi and liver fibrosis was evaluated on gross pictures of the visceral and diaphragmatic aspect of the livers after necropsy. The presence of tortuous whitish scars was scored as 1 (mild): 0-10% of the liver surface affected, 2 (moderate): 10-20% of the liver surface affected, 3 (severe): 20-30% of the liver surface affected and 4 (very severe): more than 30% of the liver surface affected. The experiment was approved by the Bioethics Committee of the University of Cordoba (code No. 1118) and conducted in accordance with European (2010/63/UE) and Spanish (RD 1201/2005) directives on animal experimentation. The number of experimental animals was set to 8 per group from a power calculation performed for a similar study conducted by our group in cattle by (18).

**Fig. 1.**
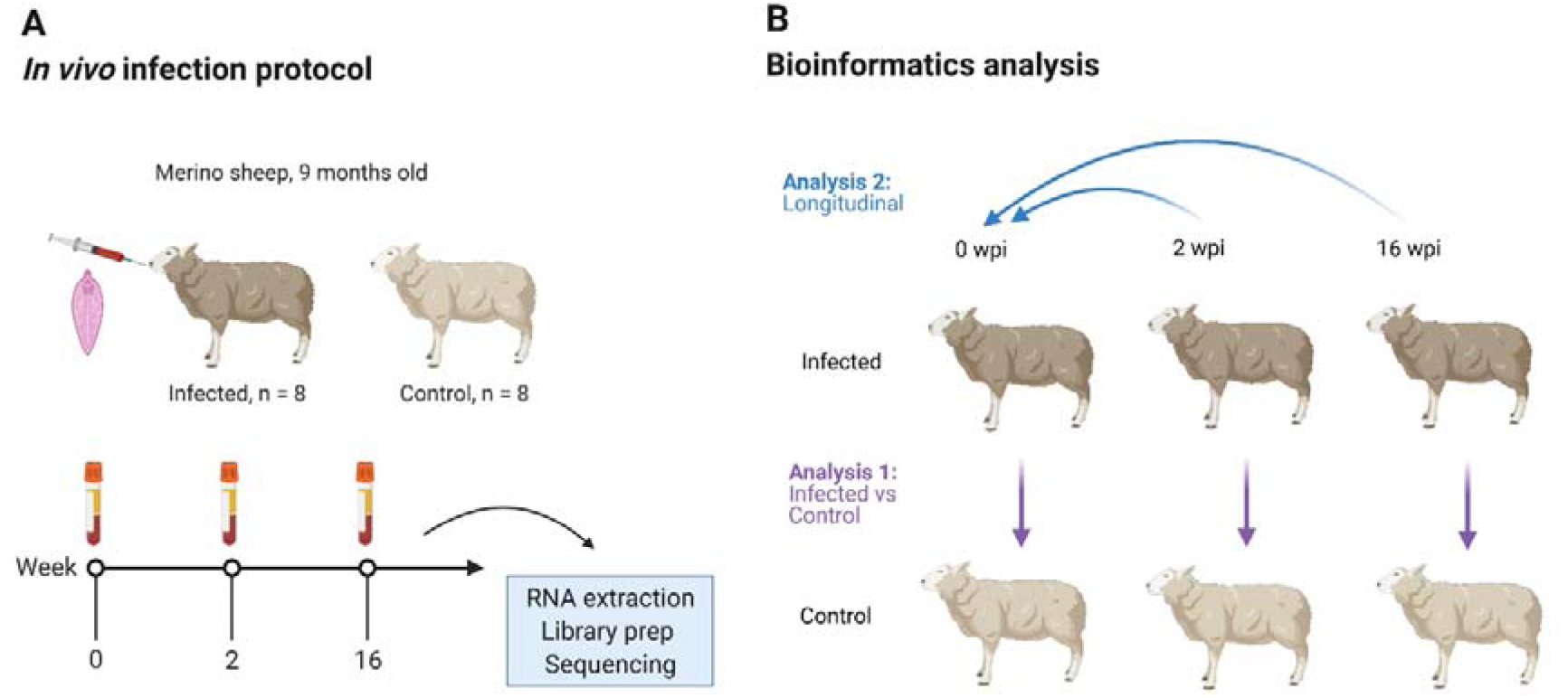
Overview of the animal infection protocol (A) and bioinformatics analysis (B). Sixteen male Merino-breed sheep were randomly assigned to an infected or control group and orally infected with a single dose of liver fluke (*Fasciola hepatica*). Blood was then collected for isolation of peripheral blood mononuclear cells (PBMC). These cells were used to generate poly-A selected, stranded 2 × 150 bp RNA sequencing data on a NovaSeq 6000, at 0, 2 and 16 wpi. These data were then analysed for DE genes using two approaches: analysis 1 (infected *vs* control) - infected animals were compared to control animals at each time point; analysis 2 (longitudinal) - the infected and control animals were compared separately to their pre-infection time points. The DE gene lists of both groups were then compared, and the DE genes which occurred in the infected group but not in the control group at each time point were considered to be the potential DE genes in response to *F. hepatica*. wpi - weeks post-infection. Image created using biorender.com

### 2.2 RNA extraction from peripheral blood mononuclear cells (PBMC)

Blood samples were collected in 10 ml EDTA vacutainers (BD) from the jugular vein. Under a flow hood, 30 ml of whole blood per animal was added into Falcon tubes and centrifuged at 1900 × *g* for 30 min without brake at 4°C to separate the buffy coat, red blood cells and plasma. The buffy coat was aspirated, added to a 15 ml Falcon tube and diluted 1:1 with PBS. After resuspension, the diluted buffy coat was gently layered on the top of 4 ml Ficoll Histopaque (Sigma-Aldrich) and centrifuged at 1900 × *g* for 30 min without brake at 4°C. Then, the buffy coat (PBMC) formed in the interphase between Histopaque and medium was aspirated. PBMC were washed once with 10 ml sterile PBS-EDTA and centrifuged at 1100 × *g* for 10 min at 4°C. After discarding the supernatant, the cells were resuspended in 5 ml of sterile ammonium chloride potassium, ACK buffer (155 mM NH_4_Cl, 10 mM KHCO_3_ and 0.1 mM ethylenediaminetetraacetic acid, EDTA) for 2 min at room temperature to lyse the erythrocytes. Then, sterile PBS-EDTA was added up to 14 ml and centrifuged at 1100 × *g* for 10 min at 4°C. The supernatant was discarded, and 10 ml of sterile PBS were added. At this step, PBMC were counted using Trypan blue for viability assessment. After counting, tubes were centrifuged at 1100 × *g* for 10 min at 4°C and PBMC were resuspended in 1.5 ml of sterile PBS, divided into cryotubes and centrifuged at 900 × *g* for 10 min at 4°C. After the PBS was removed, 1 ml of RNAlater (Sigma-Aldrich) was added per cryotube. The tubes were kept at 4°C overnight and stored at −80 °C until further assays.

Cell pellets were stored in RNAlater at −80°C until RNA extraction. Prior to extracting RNA, 950 μl of sample in RNAlater was mixed with 950 μl PBS and centrifuged at 3000 × *g* for 5 min. RLT buffer (Qiagen) was then added to the pellet and RNA was extracted using an RNeasy Plus Mini kit (Qiagen) as per the manufacturer’s instructions. RNA was eluted in 35 μl of RNAse-free water (Sigma).

RNA quality was assessed using 2100 Bioanalyzer RNA Nano chips (Agilent) as well as with a NanoDrop™ 1000 spectrophotometer (Thermo Fisher Scientific). Sample RIN values ranged between 7.3 and 9.9 (mean RIN value 9.1) and concentrations ranged from 56 – 450 ng/μl (mean RNA concentration 158 ng/μl) (Supplementary Table S1).

### 2.3 Library preparation and sequencing

RNA was sent on dry ice to Novogene (Beijing, China) for library preparation and sequencing. RNA quality was re-checked on arrival using a 2100 Bioanalyzer (Agilent) and Nanodrop. The RIN values as assessed by Novogene ranged from 7.7 to 10 (mean RIN value 9.45) (Supplementary Table S1). Stranded, poly-A selected mRNA library preparation was performed with approx. 800 ng of RNA per sample, using a NEBNext^®^ Ultra Directional RNA Library Prep Kit (New England BioLabs) according to the manufacturer’s instructions. A 5400 Bioanalyzer (Agilent) was used to assess library fragment size distribution, and qPCR was used for library quantification. The average library fragment size was 230 bp and ranged from 180 bp to 280 bp. Pooled libraries were sequenced (150 bp, paired-end) on a NovaSeq 6000 (Illumina).

### 2.4 Quality control, assembly and DE gene analysis

All the bioinformatics pipeline bash and R scripts used for computational analyses were deposited in a GitHub repository at https://github.com/DagmaraNiedziela/RNAseq_Fhepatica_ovine_PBMC. RStudio v1.3.1093 and R v4.0.3 were used for the analyses. Sequence quality was assessed using FastQC v0.11.8 (22). Trimming was performed using fastp v0.19.7 (23) with default settings as well as enabling base correction for paired end data, overrepresented sequence analysis, paired-end adapter removal, a minimum trimmed read length of 30 bases and a minimum phred score of 20. On average, 1% of the reads were trimmed per sample. Trimmed fastq files were re-checked with FastQC to confirm sequence read quality. Trimmed sequences were mapped to the ovine genome (Rambouillet v1.0) using the STAR aligner v2.7.3a (24), with gene counts generated simultaneously with the STAR software using the Ensembl annotation v101 of the *Ovis aries* reference genome and annotation. Genes with read counts < 10 across all samples as well as non-protein coding genes were removed in a filtering step (25). DeSeq2 v1.30.0 (26) was then used to visualise gene expression data and perform differentially expressed (DE) gene analysis. Data was normalised with a vst function and visualised with a principal component analysis (PCA) plot of the samples based on the 500 genes with the highest inter-sample variation. Two types of DE gene analyses were performed (**Fig. 1B**). For **Analysis 1** (Infected versus Control), DE gene analysis was performed using a negative binomial generalized linear model, with time and group as fixed effects, the individual animal included as a within-group effect, and a group by time interaction. Wald tests were performed to identify DE genes between infected and control animals at each time point, with *P* values adjusted for multiple comparisons using the Benjamini and Hochberg (B-H) method (27) DE genes with an adjusted *P* value (*P*_adj_.) < 0.1 were used for further data exploration and downstream analyses. In **Analysis 2** (Longitudinal), raw counts of both the infected and control groups post-infection were compared to their day 0 values. Animal as batch effect was not included in this model, as animals from each group were considered as their own individual controls. A difference between the gene lists from infected and control group DE genes at each time point was examined as a potential response to *F. hepatica*. The longitudinal analysis was performed for comparison purposes, as several previous transcriptomic studies have used this approach.

### 2.5 Pathway and gene correlation analysis

Significant (*P*_adj._ < 0.1, primary analysis, *P*_adj._ < 0.05, longitudinal analysis) DE genes were converted to their one-to-one human orthologs using AnnotationDbi in R, ordered by adjusted *P* value and analysed for enrichment of Gene Ontology (GO) terms, Kyoto Encyclopedia of Genes and Genomes (KEGG) and REACTOME pathways using gProfiler R package v0.7.0 (28). When a group of DE genes matched multiple pathways, the pathway that was represented by the highest number of DE genes was selected in a filtering step. Ingenuity Pathway Analysis (IPA; Qiagen) was also used to examine overrepresented pathways and predicted upstream and downstream transcriptional regulators.

### 2.6 Cell composition analysis

Transcripts per million (TPM) values were generated from raw gene counts using a custom R script. Ovine Gene IDs converted to one-to-one human orthologs were used for the analysis. To estimate the proportion of the different immune cell populations in each somatic cell sample, a quanTIseq algorithm which is based on expression of cell specific surface marker genes was used (29).

## 3 Results

### 3.1 Parasitological and Clinical Examinations

Results of the parasitological examinations have previously been described by Naranjo-Lucena et al. (2021). Briefly, no animals showed evidence of *F. hepatica* infection by faecal examination or specific antibody ELISA prior to experimental infection, and all remained clinically normal. After experimental infection, animals in the infected group showed the presence of *F. hepatica* eggs in their faeces by 9 wpi, and eggs were detected in all 8 infected animals by the end of the study. No eggs were detected in the control animals. Animals were euthanized at 16 wpi and at post-mortem examination no flukes were detected in control animals, while the mean fluke burden in infected animals was 48 ± 11 (SD) flukes per liver. At 2 wpi, infected animals showed an increased percentage of peripheral eosinophils compared to the control group. Haemoglobin concentrations were significantly lower in the infected group at 16 wpi and infected animals also exhibited a slower weight gain and enlarged hepatic lymph nodes (21). None of the uninfected sheep showed liver fibrosis. Five infected sheep showed mild hepatic fibrosis, two of them showed moderate hepatic fibrosis and one sheep showed severe hepatic fibrosis (**Supplementary Table S1**).

### 3.2 Sequence and mapping quality

PBMC collected pre-infection and at 2 and 16 wpi were used for 150 bp stranded paired-end sequencing. On average, 94% of the bases across all samples had a phred score of ≥ 30 and 98% had a score of ≥ 20. The mean number of reads per sample was 64.6 M. The distribution of read counts, phred scores and GC content per sample is available in **Supplementary Table S1**. Quality control, as assessed by FastQC, yielded satisfactory results for all samples. On average, 89.3% of reads mapped uniquely to the ovine genome Oar Rambouillet v1.0.

### 3.3 Data exploration

Raw read counts were used for DE gene analysis using DESeq2. After filtering for low counts (sum of reads from all samples < 10) and non-protein coding genes, 15,363 genes were retained for further analyses. The expression of the 500 most variable genes was used for generating PCA plots within the DESeq2 software package. Samples were observed to separate by time rather than by group, with 2 wpi and 16 wpi forming distinct clusters (**Fig. 2A**). The largest sample variation was observed at time 0, with samples dispersed across the PCA plot. This variation may reflect inter-animal differences being more pronounced at time 0 due to an acclimatisation period. For subsequent individual time points, the clearest separation between the infected and control groups was observed at 2wpi (**Fig. 2B**).

**Fig. 2.**
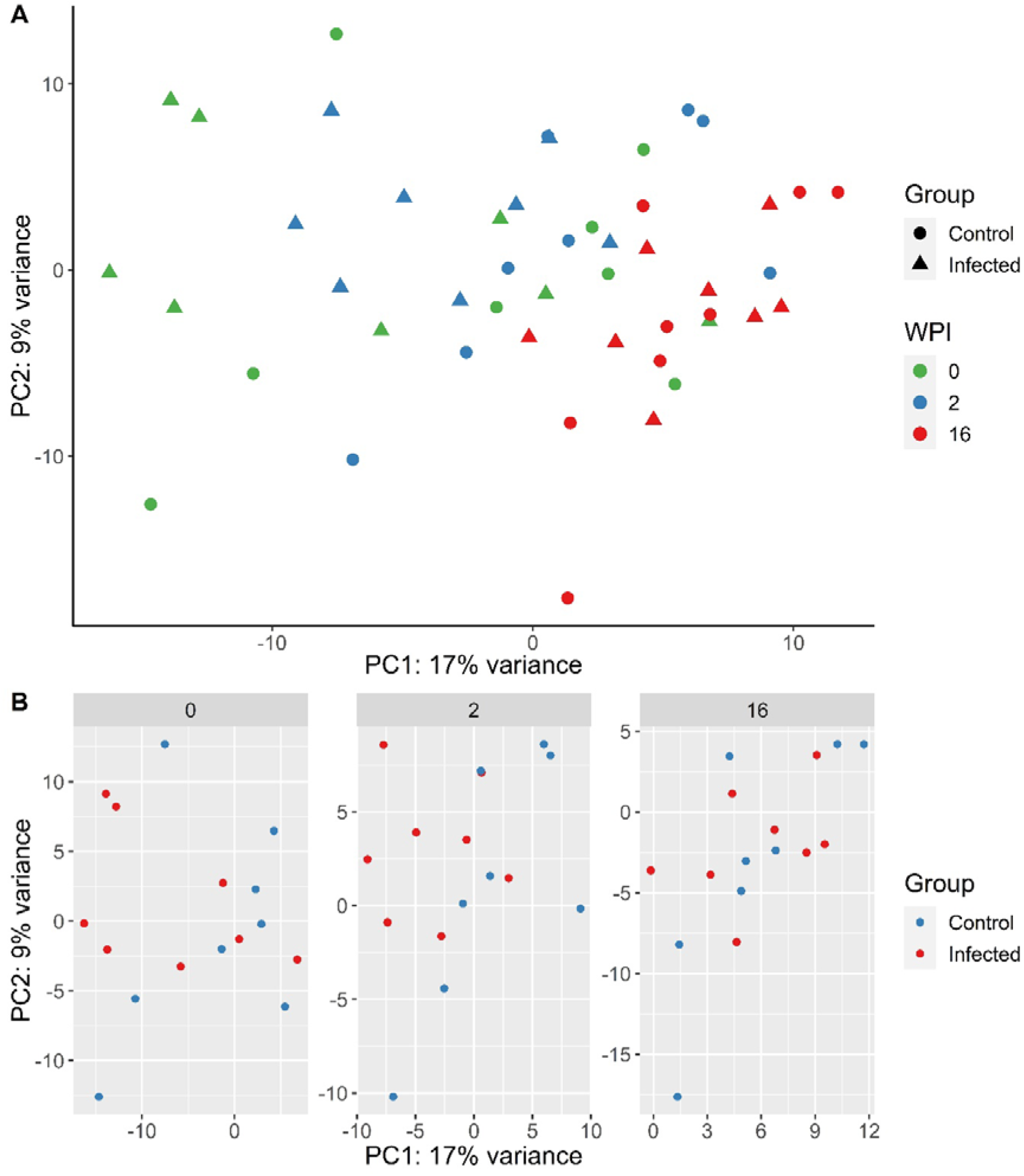
(A) PCA plot generated from PBMC gene expression data of the 500 most variable genes following infection with *F. hepatica* in the infected (circle) and control (triangle) animals. Samples from each time point are represented by a different colour: 0 wpi - green, 2 wpi - blue, 16 wpi – red. WPI - weeks post infection. (B) PCA plots generated by time point, with infected animals in red and control animals in blue.

### 3.4 Cell composition

Normalised gene expression was used to elucidate proportions of different cell types in the PBMC at each time point. There were no major differences in cell populations at any time point (**Supplementary Figure 1**), which was in accordance with haematology parameters tested previously (21). Undefined cells did not differ between groups or time points and represented 55.3 ± 0.8% in the control group and 54.8 ± 3.1% in the infected group. These “other” undefined cells most likely represent CD4 and CD8 T lymphocytes. Both *CD4* and *CD8A* were present in the data set and not differentially expressed, however the *CD8B* gene was not found in the ovine genome annotation, which was likely the reason for CD8 T-cells not being identified by QuanTISeq. Numbers of T regulatory cells appeared higher in the infected group than in the control group at 2 wpi; however, the difference was not significant. Overall, lack of major differences in cell composition indicates that the response seen in this study is likely to be due to changes in gene expression by particular PBMC cell types, rather than by changes in predominant cell type.

### 3.5 Differentially expressed genes

Analysis of infected animals versus control animals (Analysis 1) yielded 59 DE genes at 0 wpi, 453 DE genes at 2 wpi, and 2 DE genes at 16 wpi (*P*_adj._ < 0.1; **Fig. 3A**). Only 7 of the 0 wpi DE genes overlapped with the 2 wpi DE genes, and these were removed from further pathway analysis (**Fig. 3B**). The single DE gene which was common between 2 wpi and 16 wpi was *HSPB8*, which encodes the heat shock protein family B (small) member 8, and was downregulated at both time points. A complete list of infected *vs* control DE genes is included in **Supplementary Table S2**.

**Fig. 3.**
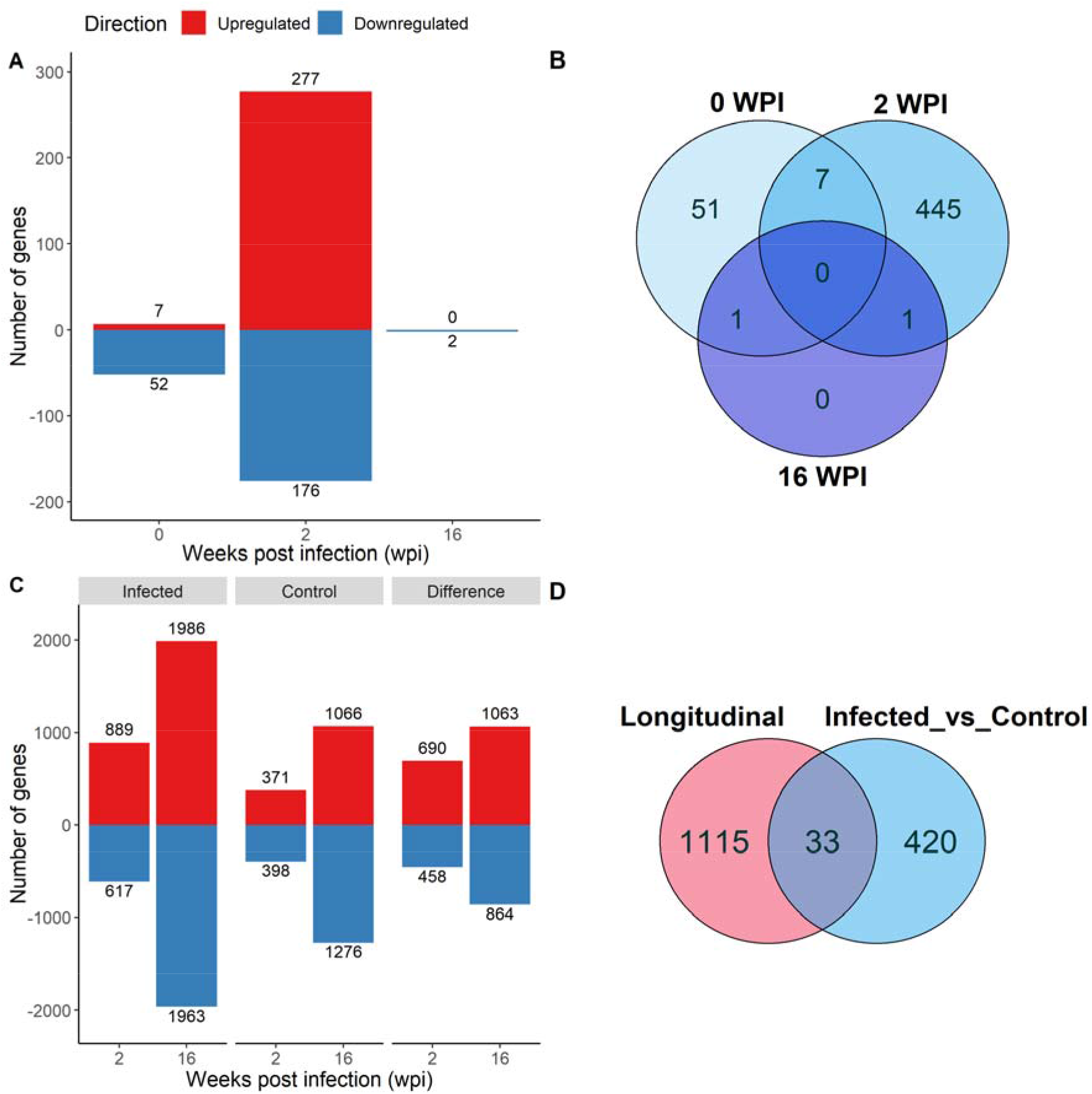
Differentially expressed (DE) genes that were up-(red) and downregulated (blue) in PBMC in response to *F. hepatica* in the iInfected *vs* Control analysis, with the Benjamini□Hochberg adjusted *P* value < 0.1 (A) and in the longitudinal analysis, adjusted P□value < 0.05 (C). Venn diagrams of overlap of genes in the different time points of Infected vs Control analysis (B) and between the Longitudinal and Infected vs Control analysis at 2 wpi (D).

To assess differential expression in both groups over time, a longitudinal analysis (Analysis 2) was performed with infected or control group compared to time 0 at each post-infection time point (2 wpi *vs* 0 wpi and 16 wpi *vs* 0 wpi in the infected group, and the same in the control group). Due to a larger number of DE genes present, a *P*_adj._ value cut-off of 0.05 was used for this analysis. A total of 1522 DE genes were detected in the infected group at 2 wpi, and 3949 DE genes at 16 wpi. In the control group there were 769 DE genes at 2 wpi and 2342 DE genes at 16 wpi. Of the DE genes identified in the infected and control groups, 1148 were found only in the infected animals at 2 wpi, and 1927 at 16 wpi (**Fig. 3C**). These 1148 and 1927 DE genes were used in further analyses. A comparison of the longitudinal Analysis 2 and the infected *vs* control Analysis 1 revealed that only 33 genes were common between the two analyses at 2 wpi (**Fig. 3D**).

In Analysis 1 (infected *vs* control), among the most significant upregulated DE genes in response to *F. hepatica*, the CD83 molecule gene (*CD83*) was a notable immune response gene in the top 5 at 2 wpi (**Fig. 4A**). CD83 is a member of the immunoglobulin (Ig) superfamily and is expressed as membrane bound or soluble forms. Membrane bound CD83 is expressed by APCs, and is most highly and stably expressed by mature dendritic cells (DCs). Soluble CD83 has been reported to have an immune suppressive function, including a role in inhibition of DCs (30). Several uncharacterised genes were in the top 5 up- and downregulated genes in the infected *vs* control analysis, as well as the longitudinal analysis (**Fig. 4B**), which may be reflective of the gaps in the functional annotation of the ovine genome.

**Fig. 4.**
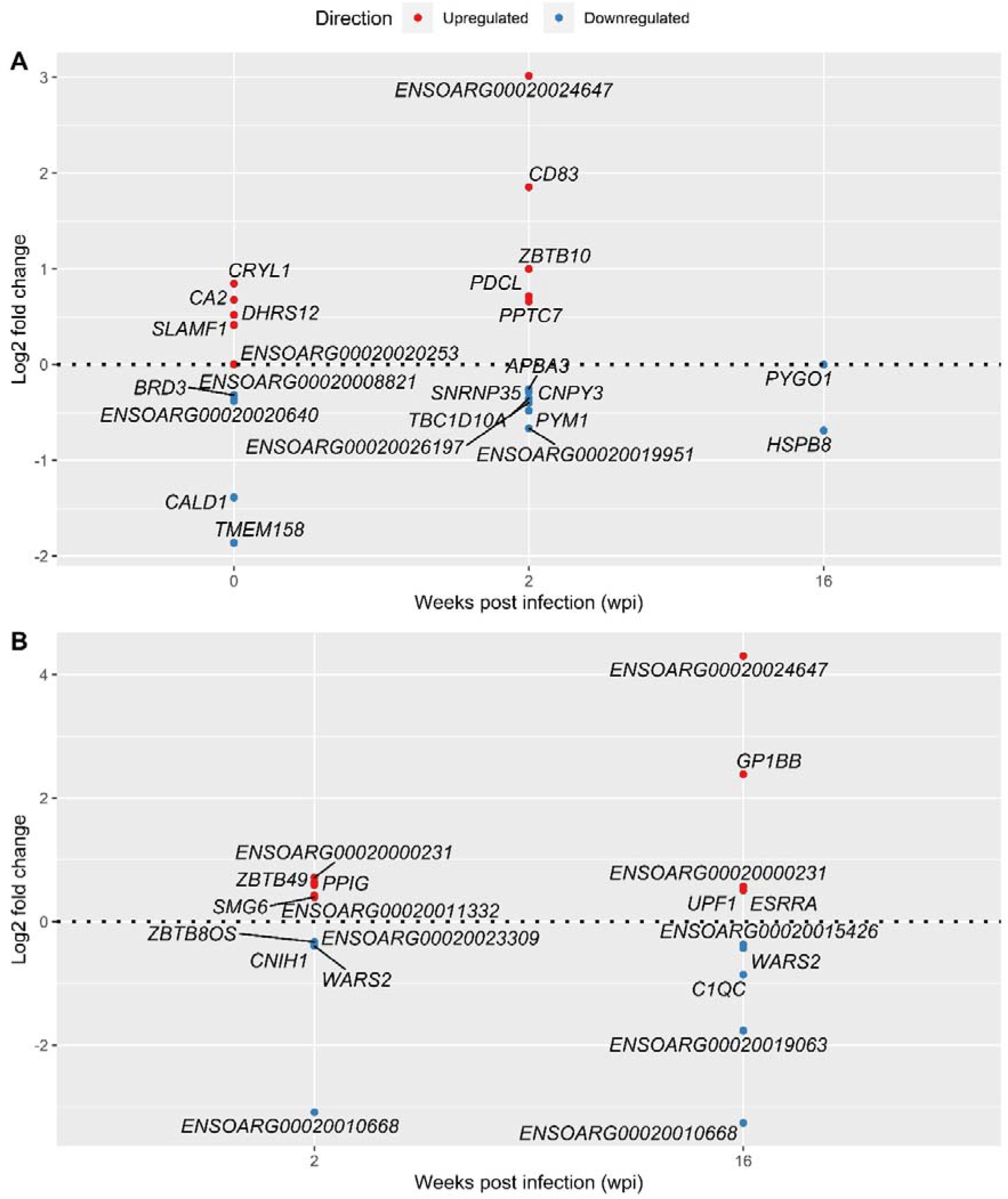
Top 5 most significant up- and downregulated DE genes in response to infection with *F. hepatica* - Infected vs Control analysis (A) and Longitudinal analysis (B). Genes with lowest adjusted *P* value were selected at each time point. In the case of genes with the same adjusted *P* value both are included.

Other notable infected *vs* control analysis DE genes at 2 wpi involved in the immune response included: the complement C1q A and B chain genes *C1QA* and *C1QB*; the CD molecule genes *CD68, CD80* and *CD151*; heat shock protein genes *HSPA13, HSPA5, HSPB1, HSPB8*; mitogen-activated protein kinase (MAPK) genes *MAP2K3, MAP3K1, MAPK11, MAPK1IP1L* as well as genes associated with TNF, NF-κB and interferon response (**Table 1**). Among these DE genes, *TGIF1* was previously found to be downregulated in cattle at 1 wpi (18) – in this study it was upregulated at 2 wpi.

**Table 1.**
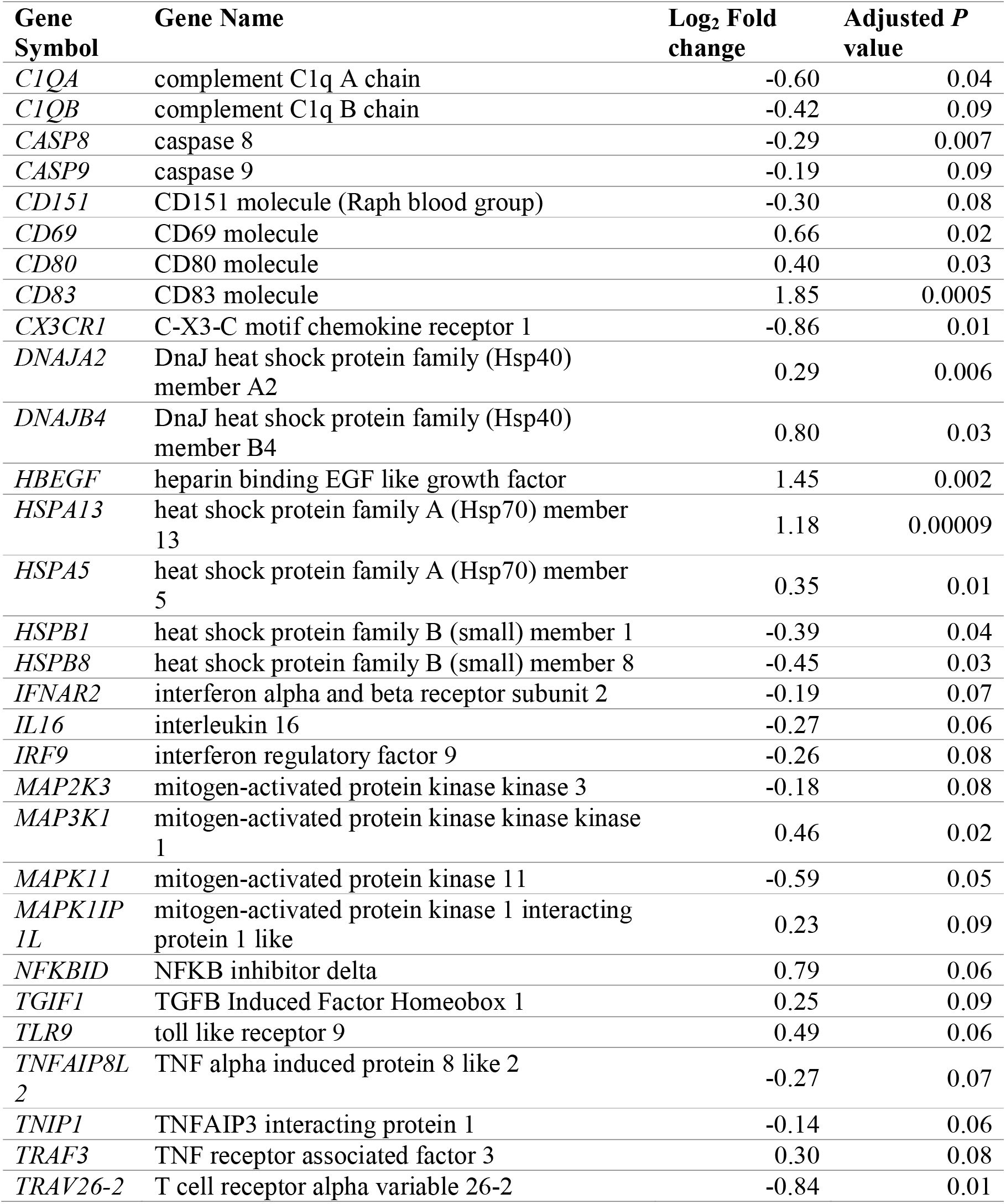
Immune response-related DE genes at 2 wpi (Infected vs Control analysis).

### 3.6 Pathway analysis of the infected *vs* control DE genes

Results of the infected *vs* control DE analysis were used to investigate overrepresented gene pathways. HUGO Gene Nomenclature Committee (HGNC) gene symbols derived from the Ensembl annotation file for the *Ovis aries* Rambouillet v1.0 genome were used for pathway analysis using IPA and gProfileR, encompassing GO, KEGG, REACTOME and IPA databases. At 2 wpi, 60 out of 453 (13.24%) DE genes had no HGNC assignment and were uncharacterised when queried using the Ensembl database. Key pathways that were overrepresented at 2 wpi included KEGG pathways such as *RIG-I-like receptor (RLR) signalling*, *Toll-like receptor (TLR) signalling* and *protein processing in endoplasmic reticulum*; and REACTOME pathways *TLR Cascades* and *Transcriptional Regulation by TP53* (**Table 2, Supplementary Table S3**). Significant IPA canonical pathways in the infected *vs* control analysis at 2 wpi included *role of PKR in interferon induction and antiviral response*, *NRF2-mediated oxidative stress response*, *p38 MAPK signalling*, *B cell receptor signalling*, *death receptor signalling* and *protein kinase A signalling* (**Table 3, Supplementary Table S4**). There was a concordance between IPA and KEGG, since *TLR signalling*, *role of RLRs in antiviral innate immunity*, and *protein ubiquitination* were also among the significant IPA canonical pathways. Furthermore, examination of GO biological processes confirmed the findings of KEGG, REACTOME and IPA, notably since pathways related to stress, protein ubiquitination, apoptosis and cell death were overrepresented (**Table 2**).

**Table 2.**
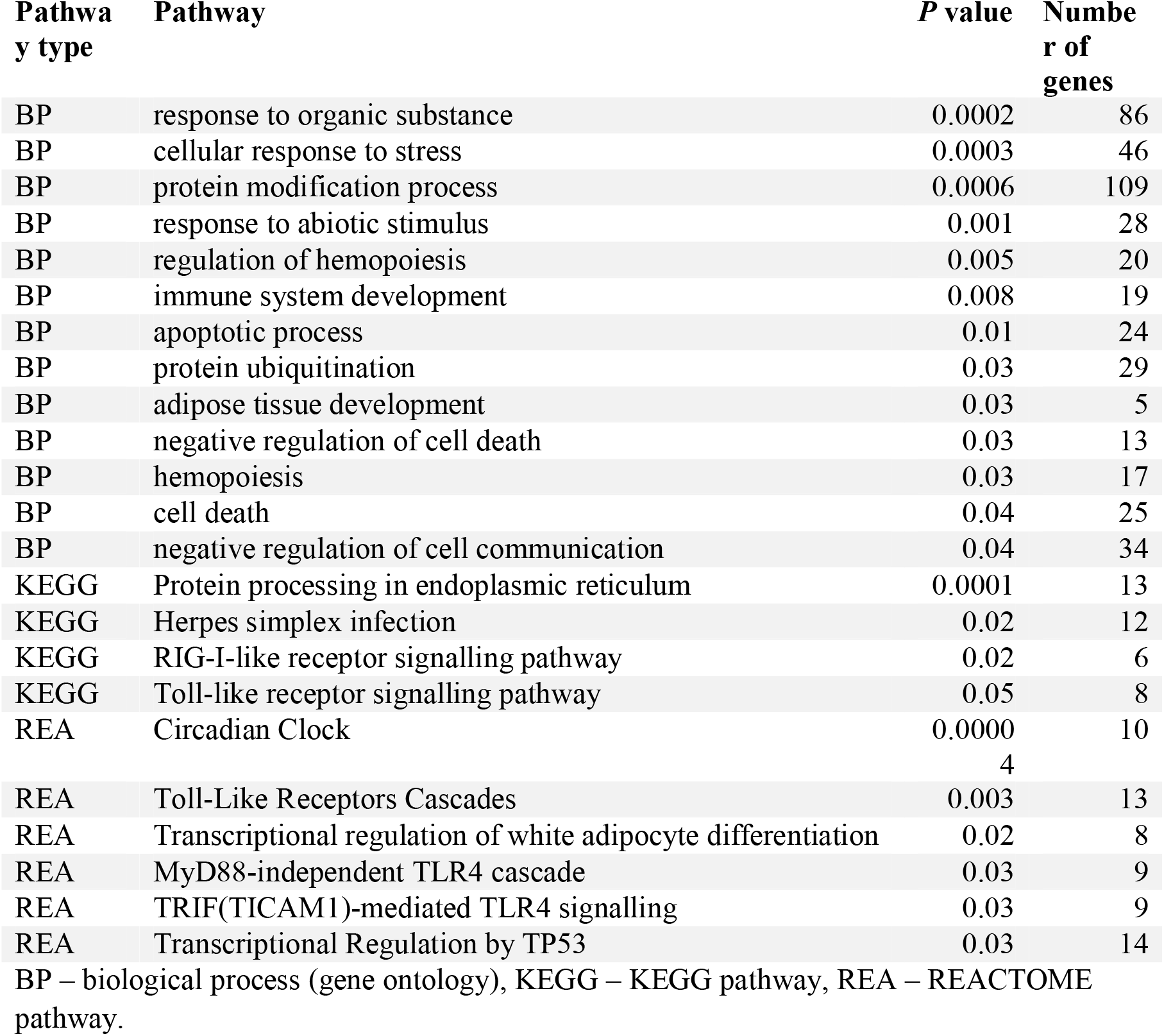
Selected pathways overrepresented in response to *F. hepatica* at 2 wpi (infected vs control analysis), as identified by gProfileR.

**Table 3.** IPA canonical pathways in PBMC at the acute stage of *F. hepatica* infection in sheep as identified by the infected vs control analysis, with a significance cut-off of p=0.01 (– log(p-value) of 2).

| Ingenuity Canonical Pathways | $-\log(P\text{ value})^a$ | Ratio<sup>b</sup> | z-score<sup>c</sup> |
| --- | --- | --- | --- |
| Role of PKR in Interferon Induction and Antiviral Response | 5.25 | 0.09 | -1.51 |
| D-myo-inositol (1,4,5,6)-Tetrakisphosphate Biosynthesis | 4.49 | 0.08 | 1.27 |
| NRF2-mediated Oxidative Stress Response | 4.00 | 0.06 | 0.71 |
| p38 MAPK Signalling | 3.72 | 0.08 | 0.71 |
| 3-phosphoinositide Degradation | 3.46 | 0.06 | 1.00 |
| B Cell Receptor Signalling | 3.44 | 0.06 | 1.00 |
| D-myo-inositol-5-phosphate Metabolism | 3.43 | 0.06 | 1.00 |
| Unfolded protein response | 3.43 | 0.11 | 1.34 |
| Endoplasmic Reticulum Stress Pathway | 3.41 | 0.19 | 1.00 |
| Protein Kinase A Signalling | 3.26 | 0.04 | -1.07 |
| 3-phosphoinositide Biosynthesis | 3.24 | 0.06 | 1.00 |
| FGF Signalling | 3.24 | 0.08 | 0.82 |
| Superpathway of Inositol Phosphate Compounds | 3.20 | 0.06 | 1.27 |
| Endocannabinoid Cancer Inhibition Pathway | 3.11 | 0.06 | -1.67 |
| Coronavirus Pathogenesis Pathway | 2.96 | 0.06 | -0.33 |
| CD27 Signalling in Lymphocytes | 2.69 | 0.09 | 1.00 |
| PEDF Signalling | 2.56 | 0.07 | 0.00 |
| TWEAK Signalling | 2.55 | 0.11 | -1.00 |
| April Mediated Signalling | 2.33 | 0.10 | NA |
| Death Receptor Signalling | 2.31 | 0.07 | -0.82 |
| B Cell Activating Factor Signalling | 2.29 | 0.10 | NA |
| ATM Signalling | 2.20 | 0.06 | 0.45 |
| Role of RIG1-like Receptors in Antiviral Innate Immunity | 2.18 | 0.09 | NA |
| Protein Ubiquitination Pathway | 2.13 | 0.04 | NA |
| ERK5 Signalling | 2.12 | 0.07 | 1.34 |
| Hypoxia Signalling in the Cardiovascular System | 2.07 | 0.07 | NA |
| Toll-like Receptor Signalling | 2.02 | 0.07 | NA |
| GNRH Signalling | 2.01 | 0.05 | -0.378 |

<sup>a</sup> $-\log(P\text{ value})$ - The probability of association of genes from our dataset with the canonical pathway by random chance alone using Fisher's exact test.
<sup>b</sup> ratio - The number of genes in our dataset that meets the cut-off criteria ( $FDR < 0.05$ and detected only in the infected group), divided by the total number of genes involved in that pathway.
<sup>c</sup> z-score - Overall activity status of the pathway. $Z < 0$ indicates a prediction of an overall decrease in activity while $Z > 0$ predicts an overall increase in the activity. NA indicates pathways that are currently ineligible for a prediction.

Key groups of genes among the significant pathways were involved in response to *F. hepatica* at 2 wpi. For instance, in the *Transcriptional Regulation by TP53* pathway, Polo-like kinase (PLK) genes *PLK2* and *PLK3* and BTG anti-proliferation factor 2 gene *BTG2* were among the most upregulated genes (**Fig. 5A**). In the *TLR signalling* pathway, the only upregulated TLR gene at 2 wpi was *TLR9*, and there was involvement of downregulated caspase gene *CASP8*, MAPK genes *MAP2K3, MAPK11*, and interferon alpha and beta receptor subunit 2 gene *IFNAR2* (**Fig. 5B**). At the same time, another MAPK gene (*MAP3K1*) was upregulated. In the *RIG-I-like receptor signalling* pathway the main antiviral receptor, retinoic acid inducible gene-I (RIG-I) encoded by *DDX58*, as well as the main transcription factor, TNF receptor associated factor 3 gene *TRAF3* were upregulated, possibly indicating pathway activation (**Fig. 5C**). On the other hand, upregulation of the NFKB inhibitor delta gene *NKFBID* indicates possible downregulation of downstream cytokine production through the NK-?B pathway. *The role of PKR in interferon induction and antiviral response* pathway involved DE genes present in the other pathways including *CASP8, DDX58, IFNAR2*, MAPK genes, *NKFBID, TLR9* and *TRAF3* (**Fig. 5D**; **Supplementary Figure 2**). *CASP9* and interferon regulatory factor 9 gene *IRF9* were also downregulated in this pathway.

**Fig. 5.**
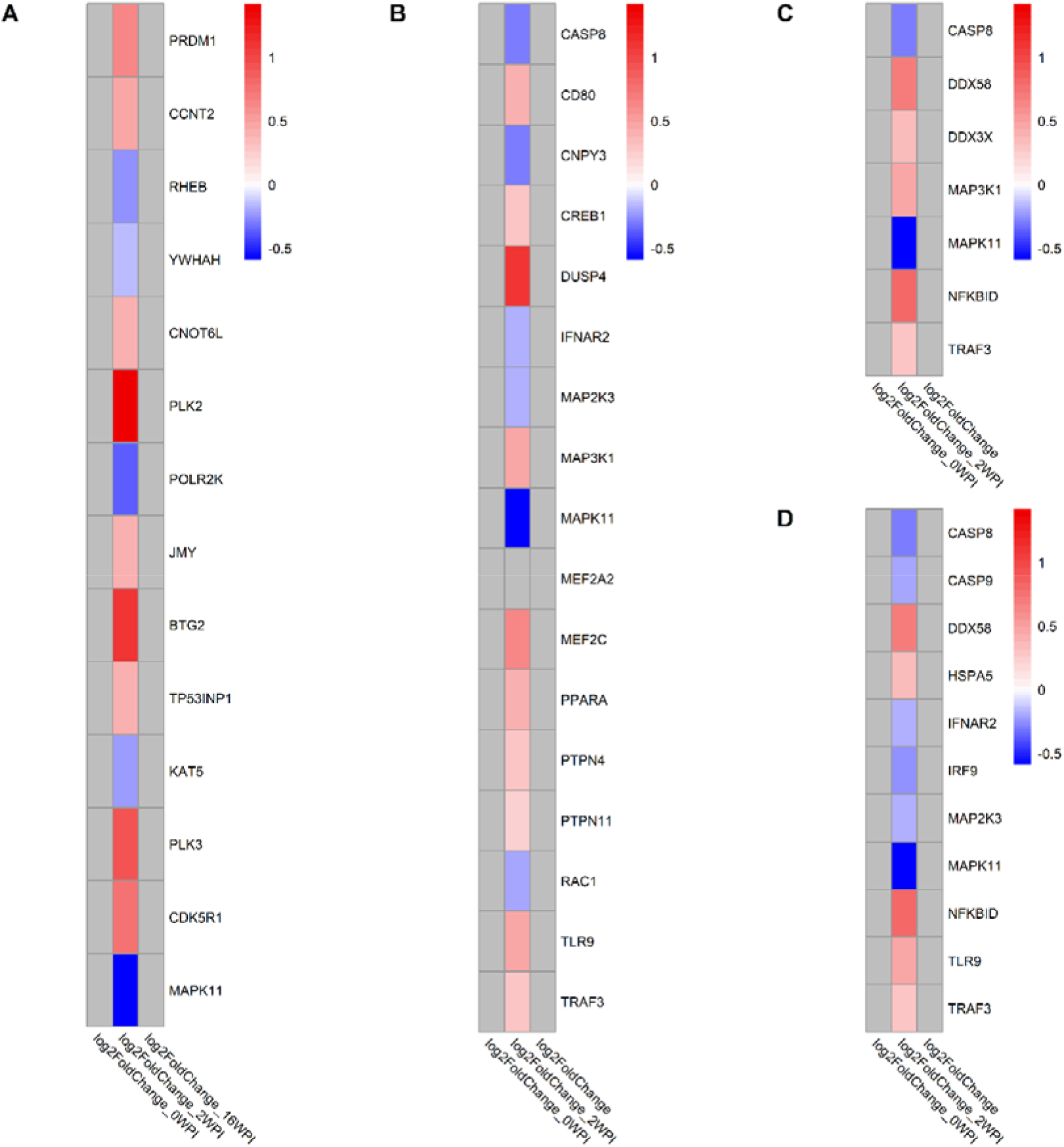
Log_2_fold change of genes involved in selected KEGG, REACTOME and IPA pathways in response to *F. hepatica* (infected vs control analysis) at each time point. (A) Transcriptional Regulation by TP53 (REACTOME), (B) Toll-like receptor signalling (REACTOME, KEGG, IPA), (C) RIG-I-like receptor signalling pathway (KEGG,IPA), (D) The role of PKR in interferon induction and antiviral response (IPA). Grey tiles show genes that were not significantly DE. If pathways were found in multiple databases, gene lists were derived as a consensus of all genes in the pathway found in all databases.

### 3.7 Comparative analysis of longitudinal bovine and ovine PBMC transcriptomic response to *F. hepatica*

A previous study by our group (18) described the bovine transcriptomic response to *F. hepatica* infection and found no robust evidence of DE genes when infected and control cattle were compared at acute and chronic timepoints post-infection. Therefore a longitudinal analysis was performed in relation to time 0, and the difference between infected and control DE gene lists were considered to be the DE genes in response to *F. hepatica*. A comparable longitudinal analysis was performed in this study, with the same number of infected and control animals in both studies (*n* = 8). To elucidate differences in responses of cattle and sheep, the longitudinal DE genes between this study and the study by (18) were compared. In the bovine study, 21 DE genes were found to differ between infected and control DE gene lists at 1 wpi, and 1624 DE genes at 14 wpi (18), while 1148 DE genes at 2 wpi, and 1927 at 16 wpi were detected in the current ovine study (**Supplementary Table S5**). The number of DE genes at the acute stage of infection in cattle was much lower than during chronic infection, while in sheep more than 1000 DE genes were detected at both the acute and chronic phases. This suggests that although in cattle penetration of immature flukes from the gut to the peritoneum elicits a relatively muted host response, the ovine response at the acute phase is more marked.

DE gene lists from the current ovine study and the bovine study were converted to human orthologs and gene lists at the acute and chronic stage were compared (**Fig. 6A**). There were three common DE genes between cattle and sheep at the acute stage of infection: *IL1RL1*, *CHD1* and *RASSF1*, and 232 common DE genes at the chronic stage of infection (**Supplementary Table S6**). When all time points in both species were compared, one DE gene was common to all time points in both sheep and cattle - the interleukin 1 receptor like 1 gene (*IL1RL1*). Common chronic DE genes included those involved in the action of TGFβ, such as the TGFB induced factor homeobox 1 (*TGIF1*) and transforming growth factor beta receptor 3 (*TGFBR3*) genes. *TGIF1* was downregulated in both species, while *TGFBR3* was upregulated in cattle and downregulated in sheep. Pro-inflammatory cytokine and chemokine genes were also among the common chronic phase DE genes, including *CCL18, CCL2*, and *IL1B*, as well as the immune receptor genes *CCRL2, IL10RA, IL2RG* and *TLR4*. Several genes were common between the species, but exhibited opposing direction of expression (**Fig. 6B**).

**Fig. 6.**
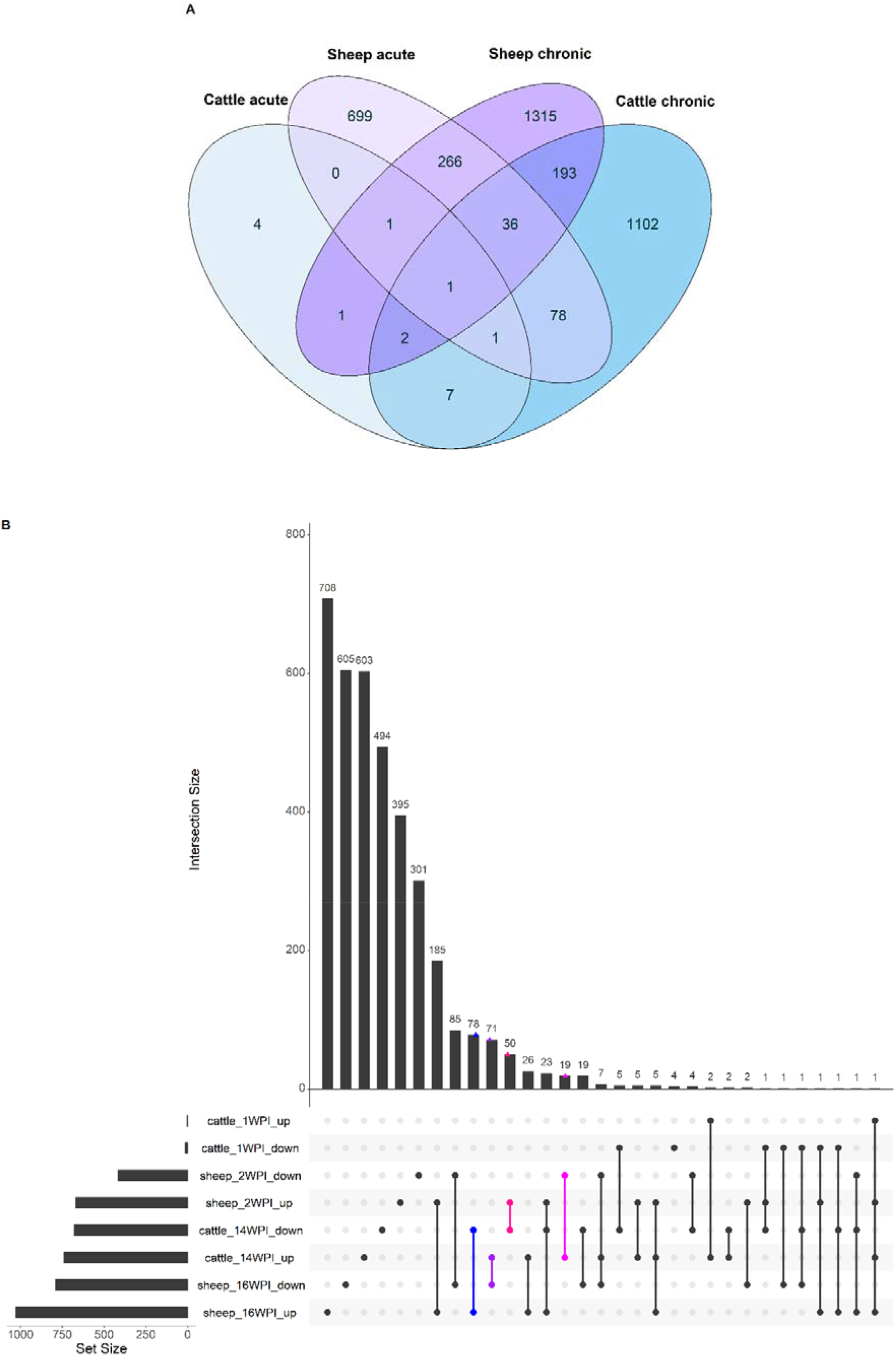
Overlap of genes from the longitudinal analysis of cattle and sheep response to infection with *F. hepatica* between the acute and the chronic phase depicted by a Venn diagram (A) and an UpSet plot where the overlaps between up- and downregulated genes are shown in both sheep and cattle at all time points (B). On the UpSet plot, overlaps where up-regulated genes are the same as downregulated genes in another species are indicated in colour (blue, purple – DE genes with opposite direction of expression in sheep and cattle chronic phase; pink, magenta - DE genes with opposite direction of expression in acute phase in sheep and chronic phase in cattle).

Pathway analysis of the longitudinal DE gene lists at acute and chronic phases was performed using IPA for both species. Three IPA canonical pathways were enriched in acute phases in both sheep and cattle, and eight canonical pathways were enriched in chronic phases for both species. The common acute pathways between sheep and cattle included *Toll-like receptor signalling*, *PPAR activation*, and *RAR activation* (**Fig. 7**). In sheep two of these common pathways were overrepresented in both acute and chronic phases of infection. *TREM1 signalling*, *STAT3 pathway* and *role of IL-17F in allergic inflammatory airway diseases* were upregulated in both species at the chronic stage. *CD28 signalling in T Helper cells*, *Phospholipase C signalling* and *NF-kB signalling* were upregulated in the chronic phase in sheep, but downregulated in cattle, while the the *hepatic fibrosis* pathway was ineligible for prediction of activation/inhibition. A total of seven pathways were significant in the chronic phase in sheep and in the acute phase in cattle, and included *IL-10 signalling, IL-6 signalling, Th2 pathway* and *Th1 and Th2 activation pathway*, which were all upregulated in sheep.

**Fig. 7.**
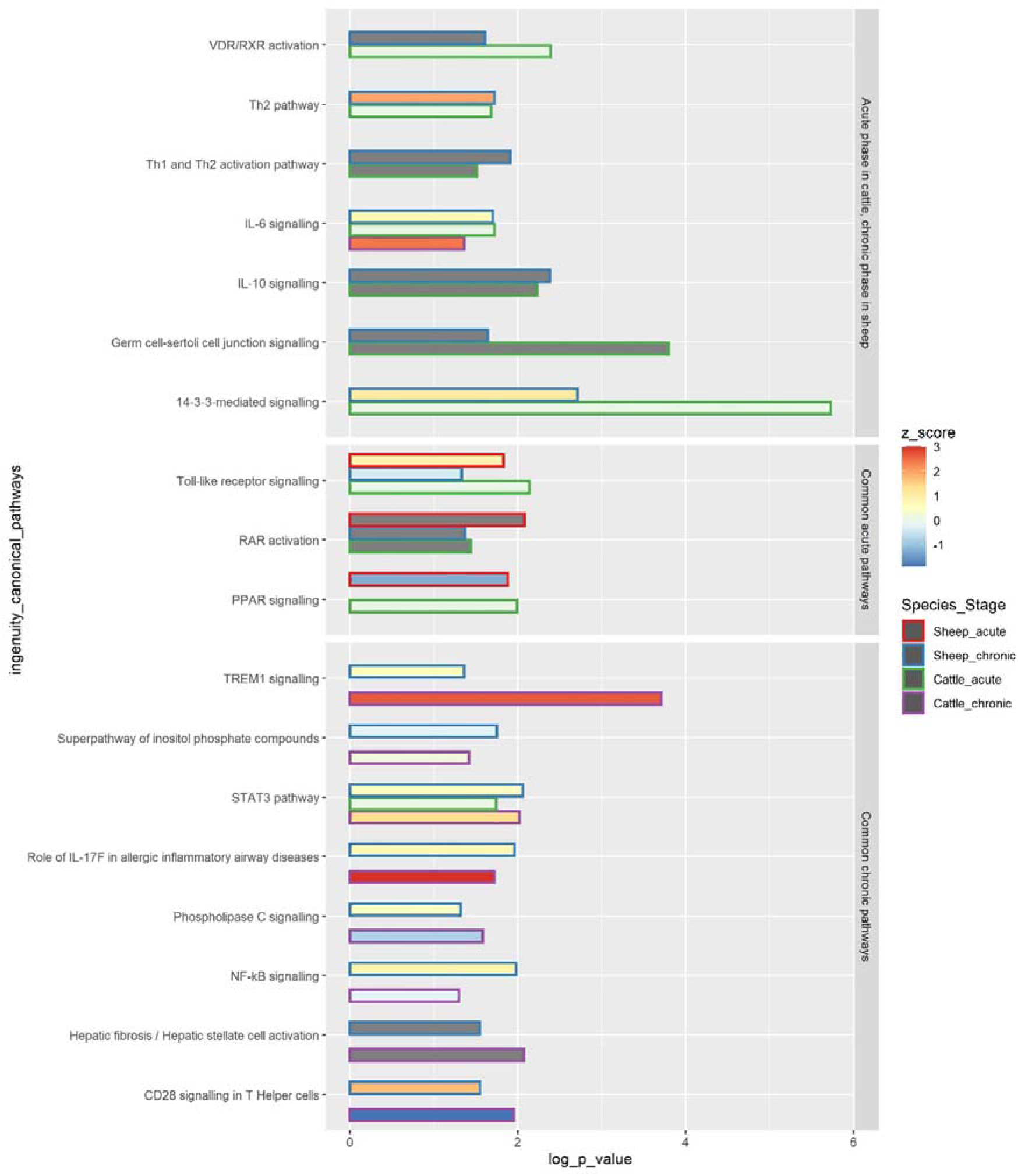
Common IPA canonical pathways in PBMC at the acute and chronic stages of *F. hepatica* infection in sheep and cattle. The common pathways are subdivided into 3 squares: pathways occurring in the acute phase in cattle and in the chronic phase in sheep, common acute pathways and common chronic pathways. The length of the bar represents -log *P* value, colour within the bar represents the z score, with the z score of 0 being represented by a light green colour, and z score not determined represented by grey bar colour. The outline of the bar represents the species (sheep or cattle) and disease stage (acute or chronic). –log(*P* value) - The probability of association of genes from our data set with the canonical pathway by random chance alone using Fisher’s exact test. z-score - Overall activity status of the pathway. Z < 0 indicates a prediction of an overall decrease in activity while Z > 0 predicts an overall increase in the activity. Grey colour of the bar indicates pathways that are currently ineligible for a prediction.

Many of the pathways specific to sheep were significant in both acute and chronic phases, and included *EIF2 signalling, NGF signalling, NER Pathway, SAPK/JNK signalling* and *B Cell Receptor signalling* (**Supplementary Table S7**).

## 4 Discussion

*F. hepatica* is an important pathogen that causes zoonotic disease and economic losses in animal agriculture. Due to lack of acquired immunity following natural infection (31; 32) and anthelmintic resistance concerns, there is an urgent need for the development of novel control methods such as vaccination. Over the last few decades, progress has been made in the isolation, characterisation and testing of a number of native and recombinant molecules as vaccines against liver fluke disease in ruminant hosts (33). However, data from vaccine trials have been inconsistent (14), and some recent trials with recombinant antigens failed to induce protection against fluke infection (20) or achieved only a minor reduction of fluke burdens (34). In spite of evidence that effective vaccination may be possible, design of smart vaccines will require more knowledge about the immune response to *F. hepatica*, which can be provided by big-data approaches such as transcriptomics. This study provides an insight into the response of ovine PBMC to *F. hepatica* at both the acute and chronic stages of infection, and is the first long-term ovine PBMC transcriptomic study that includes uninfected control animals in the experimental design.

### 4.1 Infected *vs* control analysis

#### 4.1.1 RIG-I like and Toll-Like receptor signalling pathways are identified by multiple databases and indicate potential production of interferons and downregulation of apoptosis

Two databases (KEGG and IPA) identified the RLR signalling pathway as significant, and TLR signalling was significantly overrepresented in 3 databases (KEGG, REACTOME and IPA).

##### 4.11.1 RIG-I like receptor signalling

RLRs are cytosolic RNA helicases that recognise viral RNA inside immune and non-immune cells, leading to recruitment of intracellular adaptor proteins and ultimately to production of type I interferon and inflammatory cytokines. RLRs include RIG-I, melanoma differentiation-associated gene 5 (MDA5), and DExH-box helicase 58 (DHX58 or LGP2) (35). The key receptor in this pathway, RIG-I, is encoded by the *DDX58* gene, which was upregulated in this study at 2 wpi (**Fig. 5C**). RIG-I is responsible for the innate recognition of uncapped 5′-phosphate double-stranded or single-stranded RNA found in numerous viruses including influenza and Ebola (35). Another key gene that was upregulated in our study and encodes an important transcription factor is the *TRAF3* gene. Since TRAF3 is key to intracellular responses, molecules secreted by the parasite may undergo endocytosis and be detected while inside the cell. Therefore, the RLR signalling pathway could be involved in recognition of and response to extracellular vesicles of *F. hepatica*, which can contain RNA and miRNA and are secreted throughout the life cycle of the parasite, including newly excysted juveniles and adults (36; 37). TRAF3 activity in RLR signalling is further regulated by the action of Polo-like kinases (38), which were also upregulated at 2 wpi (**Fig. 5A**). Overall, the upregulation of these key genes in the RLR signalling pathway would ultimately lead to an increase in interferon production. While no interferon genes were detected as upregulated in this study, the activation may occur at a later stage in the infection.

##### 4.11.2 Toll-like receptor signalling

TLR signalling was identified using both the KEGG and REACTOME pathway databases. The only TLR gene that was upregulated in this pathway was *TLR9* (**Fig. 5B**), which is activated by unmethylated CpG sequences in DNA molecules due to cancer, infection, or tissue damage. TLR9 is the receptor involved in the response to oncogenic viruses, and in autoimmune disorders. The overexpression of *TLR9* coinciding with overexpression of *DDX58* (RIG-I) further suggests that a response to nucleic acids secreted by *F. hepatica* is taking place. TLR7 and TLR9 are selectively expressed by plasmacytoid dendritic cells (pDCs; also known as IFN-producing cells), which are a subset of DCs with a plasmacytoid morphology and unique in their capacity to rapidly secrete large quantities of type I IFN in response to viral infection, including response to influenza (39). Glycans produced by *F. hepatica* were previously reported to induce a semi-mature state of DCs in mice, with reduced IFN-γ production (40). Excretory-secretory products (ESP) from *F. hepatica* have also been found to inhibit the production of effector molecules such as IL-6 and IL-12p70 in DCs following stimulation of TLR9 with CpG (41). The DC receptor genes *CD80* and *CD83* were among the most upregulated DE genes in this study at 2 wpi (**Table 1, Fig. 4A**) – which may further suggest initial activation of DCs in response to *F. hepatica*, with further downstream response being inhibited by the parasite.

The lack of differential expression of other TLR genes in this study is in contrast to a previous study of transcriptomic response of ovine PBMC conducted by our group, where *TLR1, TLR5, TLR6, TLR7* and *TLR10* were downregulated in PBMC at the acute stage of infection (20). However, in a separate study of ovine PBMC, no TLR genes were DE at 2 wpi (19). TLR signalling also involves upregulation of TRAF3, which undergoes endocytosis together with TLR4 in response to lipopolysaccharide (LPS). LPS has been identified as a predicted upstream regulator by IPA at 2 wpi (results not shown), which may indicate a potential influence of helminth-derived LPS (perhaps from endosymbiotic bacteria) on the ovine response. Various other helminths have been demonstrated to possess gut endosymbionts that are transferred across generations and not present in the host’s gut, including *Neorickettsia* in *Plagiorchis elegans* and *Weissella* and *Leuconostoc* in *Haemonchus contortus* (42). *Neorickettsia* has also been found in *F. hepatica* obtained from ovine liver (43). Helminth microbiomes could play a role in developmental variation and evolutionary fitness of the parasites and may influence the host immune response.

Ultimately, the TLR signalling pathway leads to upregulation of NF-kB and MAPK related genes. In this study, the NF-kB inhibitor delta gene *NFKBID* was upregulated, and MAP kinases downregulated (**Fig. 5B**), both of which would lead to a decreased cytokine response. In fact, we did not detect any interleukin genes to be DE apart from *IL16*, which points to an inhibition of cytokine response taking place.

Caspase 8 is a common protein in both RIG-I and TLR signalling pathways and is known to be a key driver of cellular apoptosis (44). This caspase was among the most downregulated DE genes at 2 wpi, and suggests a potential inhibition of cellular apoptosistaking place at the acute stage of infection. Caspase 8 also may be key to the inflammatory response as *casp8*^-/-^ mice exhibited a reduction in *IL1B* and *IL12B* expression, reduced recruitment of monocytes and an increased susceptibility to *Toxoplasma gondii* (45).

#### 4.1.2 Several pathways related to apoptosis and cell death were identified

##### 4.1.2.1 Transcriptional regulation by TP53 was overrepresented

*Transcriptional Regulation by TP53* is a REACTOME pathway that groups the upstream and downstream effects of the transcriptional regulator p53, most known for being involved in cancer regulation. The transcription factor recognizes specific responsive DNA elements and regulates the transcription of genes involved in cellular metabolism, survival, senescence, apoptosis and the DNA damage response (46; 47). *PLK2, PLK3* and *BTG2* were the most upregulated genes in the pathway (**Fig. 5A**), encoding the PLK2 and 3 and BTG anti-proliferation factor 2 proteins, respectively. PLKs play pivotal roles in cell cycle progression. PLK2 and 3 are activated by spindle checkpoints and DNA damage, and are required for entry into S phase (48). The fact that the *PLK2* and *PLK3* genes are highly upregulated here would indicate a progression of the cell cycle and activation of cell proliferation. *BTG2* belongs to the anti-proliferative (APRO) family of genes that regulate cell cycle progression by enhancing or inhibiting the activity of transcription factors. The *BTG2* gene has a potential role in muscle fibre size, intramuscular fat deposition and weight loss in sheep and can lead to a decrease in cell proliferation or an increase in energy expenditure (49). Therefore, upregulation of this gene may be related to increased cellular metabolism.

Other DE genes at 2 wpi that may indicate perturbation of the cell cycle include the cyclin D3, L1, Q and T2 genes (*CCND3, CCNL1, CCNQ, CCNT2*) and cyclin-dependent kinase genes *CDK17* and *CDK5R1*, with the majority of these being upregulated (**Supplementary Table S2**). The product of the *CCND3* gene, downregulated in this study, is responsible for the progression through the G1 phase (50). Cyclin types L and T, which were upregulated in this study, are involved in transcription (51; 52). It is therefore likely that transcription is activated at 2 wpi rather than PBMC proliferation.

##### 4.1.2.2. The role of PKR in interferon induction and antiviral response pathway was inhibited at 2 wpi

Protein kinase receptor (PKR) belongs to a family of pattern-recognition receptors (PRRs). It can be induced by interferons (IFNs) as well as dsRNA from viral sources and responds by stimulating apoptosis of the host cell or by inhibition of translation, affecting both viral and cellular mRNA. Apoptosis can be activated via death receptor pathways such as caspases, as well as by direct phosphorylation of p53. Much of PKR signal transduction, however, occurs via NF-?B and MAPK pathways, which trigger various antiviral effects especially the production of type I IFNs and increased expression of pro-apoptotic factors (53). The interferon alpha and beta receptor subunit 2 (*IFNAR2*) and interferon regulatory factor 9 (*IRF9*) genes were downregulated in this pathway (**Supplementary Figure 2**). IRF9 is a transcription factor which mediates signalling by type I IFNs (IFN-α and IFN-β). Following type I IFN binding to cell surface receptors such as IFNAR, STAT1 and STAT2 are phosphorylated and IRF9 associates with the phosphorylated STAT1:STAT2 dimer to form a complex termed ISGF3 transcription factor, which then enters the nucleus and activates the transcription of interferon stimulated genes thereby driving the antiviral response of the cell (54). Downregulation of these genes at 2 wpi, indicates that the interferon response is inhibited. The NFKB inhibitor delta gene (*NFKBID*) is upregulated, which suggests inhibition of the PKR signal transduction that occurs through the NF-κB pathway. MAPK genes are downregulated, which also suggests inhibition of the PKR signal transduction which occurs through the MAPK pathway. Overall, the downregulation of this pathway suggests that an inhibition of innate immune response and cellular apoptosis are taking place. These mechanisms may facilitate host defence against *F. hepatica*, but be inhibited due to the parasite’s immunomodulatory properties (55). Notably, anti-viral/small RNA response activation is suggested by upregulated genes in the TLR and RIG-I signalling pathways. This leads to the hypothesis that downstream interferon response is actively inhibited, even though foreign nucleic acids are being detected inside the cell.

##### 4.1.2.3 Death receptor signalling was inhibited at 2 wpi

Death receptor signalling and apoptosis signalling were observed to be upregulated in the acute stage of *F. hepatica* infection in a study we conducted in sheep previously (20). In this study, the IPA *death receptor signalling* pathway was inhibited. Caspase-8 is a major initiator in the death signalling pathway (44). Previously, it has been shown that *F. hepatica* ESPs induced apoptosis of eosinophils, and caspase 3, 8 and 9 were activated in the process (16). *CASP8* and *CASP9* were both downregulated in this study, and *CASP8* downregulation has also been described previously in sheep (20). Overall, the death receptor signalling pathway was inhibited; however, the key genes involved in the pathway such as *TNF* and *CASP3* were not DE. This pathway should therefore be studied in more detail in order to elucidate the mechanisms of cell survival or apoptosis during infection with *F. hepatica*.

Overall, the pathways found at 2 wpi in the infected *vs* control analysis suggest that immune response to exogenous nucleic acids as well as changes in energy metabolism and cell viability occurred in the acute phase of ovine *F. hepatica* infection. Involvement of p53 related proteins, and a RIG-I like and TLR9 mediated anti-viral response are among the new response characteristics identified in this study.

### 4.2. Longitudinal analysis

#### 4.2.1 Th2 cell activation

In cattle, a shift towards Th2 responses highlighted by involvement of Th2 response and IL-4 and IL-6 signalling pathways occurred in the acute phase of infection (18). It seems that the involvement of a Th2 response and anti-inflammatory pathways involving IL-10 occurs only in the chronic phase in sheep, while in cattle these responses are activated much earlier. Activation of Th2 cells and inhibition of Th1 cells and a corresponding shift towards anti-inflammatory pathways was also observed in a study we conducted on ovine hepatic lymph node (HLN) at 16 wpi, confirming that the anti-inflammatory responses are activated in sheep in the chronic phase of *F. hepatica* infection (21). This may suggest that it is the early activation of an anti-inflammatory response that causes lack of clinical signs in the acute infection in cattle, and that severe acute clinical signs in sheep are mainly caused by an acute inflammatory reaction in the attempt to remove the pathogen, with a corresponding lack of anti-inflammatory response to mediate tissue damage. A tissue repair type of response seems effective in preventing severe disease in cattle, but not in providing protection against re-infection (56; 57).

#### 4.2.2 TLR signalling

The TLR pathway was upregulated in sheep in the acute phase and downregulated in the chronic phase of infection, as indicated by the IPA z score (Fig. 7, Supplementary Table S7). On the other hand, TLR signalling was overrepresented (with no consensus on activation or inhibition, z score = 0) only at the acute stage in cattle (Fig. 7). In cattle, the *TLR2* and *TLR4* genes were upregulated in the chronic phase, and *TLR10* was downregulated (even though the pathway was not significant at the time), and no TLR genes were DE in the acute phase (18). On the other hand, in sheep, *TLR4* was downregulated in the chronic phase, and no TLR genes were DE in the acute phase, with *IL1RL1, LY96*, and *MAP4K4* being the main genes contributing to the activation of this pathway in both stages. TLR2 recognizes a wide variety of PAMPs including lipoproteins, peptidoglycans, lipotechoic acids, zymosan, mannan, and tGPI-mucin, while TLR4 recognizes LPS, and TLR10 has been found to recognise ligands from *Listeria* and influenza A infections (58).

#### 4.2.3 Hepatic fibrosis

The IPA *hepatic fibrosis* pathway was overrepresented at the chronic stages in both sheep and cattle (Fig. 7). TGF-β1 is an essential cytokine promoting collagen production and fibrosis, which then leads to encapsulation of flukes and limiting migration of the parasites through the liver parenchyma (11; 59). Liver fibrosis in ruminants has previously been associated with expression of *IL10* and *TGFB*, with an increased expression of these genes potentially leading to increased fibrosis and control of fluke burdens (60). In cattle, liver fibrosis has been proposed as a downstream effect of upregulated *TNF* and *IL1B*, which would activate the genes that were also upregulated in the bovine study, *i.e.*, *IL6, PLAU, SERPINE1, TNFRSF1A, SOCS1*, and *CTSB* (18). In sheep, however, the pathway was observed to exhibit a more recognisable outcome, with the involvement of *IFNG*, the collagen genes *COL11A1* and *COL11A2*, and matrix metalloprotease *MMP9* and *SMAD2* genes (**Supplementary Table S7**). Even though typically collagen type I and III are involved in chronic liver fibrosis, the upregulation of genes encoding alpha and beta subunits of collagen type XI as well as a matrix metalloprotease gene, which is involved in reorganisation of the extracellular matrix, supports a potential role of these proteins in fibrosis in chronic liver fluke infection in sheep. Interferons alpha and gamma, typically secreted by NK cells, are also thought to inhibit liver fibrosis (61; 62). *IFNG* was downregulated in the chronic phase in sheep, which further supports activation of fibrosis. Surprisingly, *SMAD2* was downregulated (**Supplementary Table S6**), which could suggest inhibition of fibrosis, even though other genes in the pathway were activated. *IL1B* and *TLR4* were involved in the pathway in both sheep and cattle, so there could also be an role in liver fibrosis activation for *IL1B*. *TNF* was not DE in sheep. The *SERPINE1 gene*, which was detected at chronic stages in the bovine study (18) and in a previous study of ovine PBMC response following infection with *F. hepatica* (20), was also not DE in this study.

The low number of DE genes in the acute response of cattle, coupled with specific anti-inflammatory pathways such as Th2 activation or IL-10 signalling, which were upregulated during the chronic phase in sheep and that were upregulated in the acute phase in cattle, underscore the significance of the acute response in sheep and its potential detrimental effect on disease phenotype. It is important to note that in sheep, following experimental infection, flukes mature simultaneously. Therefore, sheep typically yield adult flukes of approximately the same age and size in the liver during pathological examination at the chronic stage, while in cattle the chronic infection presents with a mix of juvenile and adult flukes. The more gradual progression of flukes to maturity in cattle suggests that the interaction between the parasite and the host may cause a delay in fluke maturation, potentially by inhibiting the growth of neoblasts (63). On the other hand, sheep do not seem to be able to slow the life cycle of the fluke, which then causes a rapid migration of the flukes through the gut and maturation in the liver causing a large inflammatory response, coupled with an attempt to repair tissue damage and slow inflammation through Th2 responses. The conclusion, therefore, would be that an early inflammatory response to *F. hepatica*, as seen in sheep (11; 64; 65), has major pathological consequences.

#### 4.2.4 Comparison with a previous ovine PBMC study

(18) discussed their findings in comparison with a previous transcriptomic study of ovine PBMC by (20) and concluded that in sheep, the majority of DE genes were present in the acute phase when compared to the chronic phase, while in cattle only 5% of the total DE genes were expressed in the acute phase of infection. While the proportion of acute DE genes in the bovine response is indeed very low, our study shows that using the same longitudinal analysis as in the cattle transcriptomic study, with the inclusion of control animals in the analysis, leads to a result where 63% of all DE genes were allocated to the chronic phase. Even when examining the infected animal group only, as in the study by (20), the majority of DE genes in our study were expressed during the chronic phase of infection, with 72% of the DE genes detected at this stage. It is important to note, however, that in the previous ovine transcriptomic study by (20) the DE genes at the chronic phase were compared to the acute phase (*i.e.*, T3 *vs* T2), not to the baseline of week 0. In the bovine DE gene analysis, the number of DE genes from the chronic phase compared to acute phase (WK14 *vs* WK1) was three-fold lower than when the chronic phase was compared to baseline (18). Therefore, the choice of time point comparison in the DE gene analysis could explain the skew towards the acute phase in the previous study of the transcriptomic response in sheep. Breed differences, as well as a differences in the *F. hepatica* strain used could also partially account for the different results when comparing our study to (20).

In both our study and the bovine study by (18) the log_2_ fold changes of DE genes were relatively low in comparison to the previous ovine transcriptomic study (20): in our study the longitudinal DE gene log_2_ fold changes ranged from −3.1 to 3.3 in the acute phase, and from −3.2 to 4.3 in the chronic phase. While these log_2_ fold changes are still higher than in the bovine study, the differences are relatively minor, especially given the different DE gene analysis packages used (edgeR in the bovine study *vs* DEseq2 in the current ovine study), which could lead to slightly different results (26). The previous ovine study (20) observed much larger log_2_ fold changes, ranging from −27 to 12. The study used a Limma-Voom package for DE gene analysis, which could at least partially account for the differences in log_2_ fold changes. The DE gene analysis in Limma is based on a linear model, with gene counts typically log-transformed, while DESeq2 and edgeR use a negative binomial generalised linear model, with raw counts used for the analysis. A difference in normalisation of raw counts prior to DE gene analysis could also cause differences in log_2_ fold changes.

### 4.3 Comparison with previous ovine transcriptomics studies

Our study confirmed the need to use control animals in the infection. A greater number of DE genes were identified in sheep in response to *F. hepatica* by (20). However, this previous study did not use uninfected control animals in its design, and DE genes were compared to pre-infection. In fact, when the DE gene analysis was performed in this study in comparison to time 0 (longitudinal analysis), a large number of DE genes was identified in both infected and control animals, a phenomenon also found in a previous bovine study (18). This suggests that presence of DE genes related to developmental changes that occur in 9 month sheep could obscure the response to infection. Therefore, a study on older animals may be warranted, where growth effects may not be so dominant. Another possible explanation may be the seasonality of the infection and the onset of low temperatures. A previous study by (66) identified DE genes in sheep due to wool production between the months of August and December. The resulting DE genes included *IL1B, IL5, IL8* and *CXCL1*, which suggests that immune proteins may be produced or inhibited due to seasonal changes. Our study was conducted between September and January, which could suggest a similar seasonal effect. In addition, immune response in sheep could be breed-dependent, as demonstrated by relative resistance of the Indonesian thin tail sheep breed against *F. gigantica* when compared to Merino sheep (67; 68). Our study was conducted on Merino sheep, while the previous ovine study (20) used the Suffolk breed. Longitudinal studies of gene expression in sheep with seasonal changes, potentially in conjunction with measuring hormones and other physiological variables could assist future transcriptomic studies of host response.

In a previous ovine PBMC study by (19), genes encoding for galectin 9, SMAD2 and TGF-β were all upregulated at 2 wpi. The genes and pathways described in (19) differed from genes and pathways found in this study at 2 wpi. While *SMAD2* was DE in the longitudinal analysis in this study at 16 wpi, it was downregulated. Lack of transcription of interleukin genes was also observed in the previous study (19), while in our study *IL16* was downregulated. Since both studies were performed on Merino breed sheep, with the inclusion of control animals, the differences could be attributed to a different strain of *F. hepatica* used, or seasonal differences, if the previous study was conducted in different months. Both (20) and (19) used metacercariae from Baldwin Aquatics, Inc, while in this study Ridgeway metacercariae were used. It is also notable that the DE gene analysis in (19) was performed using NOISeq, which has been shown to produce a comparably large number of false positive results when less than six biological replicates are used (69). Overall, the PBMC studies in sheep show varied results, likely due to breed and infecting strain differences, as well as due to the bioinformatics methods used. It is important to note, that while we conducted this study in PBMC in order to be able to compare to previous studies, in the future studies on whole blood immediately preserved in PAXgene (Qiagen) or Tempus (Thermo Fisher Scientific) tubes could be performed to validate the results of these ovine transcriptomic studies, as preserving whole blood immediately is thought to minimise *ex vivo* variation and RNA degradation (70).

The animals used in our study have also been used to analyse the response to *F. hepatica* in the HLN at 16 wpi (21). There were common pathways found in PBMC in the infected *vs* control analysis in this study at 2 wpi and in the HLN at 16 wpi. Specifically, B cell receptor signalling was upregulated in both tissues, and apoptosis and NK cell signalling were downregulated. Crosstalk between DCs and NK cells was downregulated in the hepatic lymph node, while in this study the direction of pathway activation could not be determined, mainly due to a low number of DE genes involved (**Supplementary Table S4**). In our study, the CD-NK pathway was mainly significant due to the activation of DC receptor genes *CD69, CD80* and *CD83*. It is important to note that the comparison is made between the acute phase in PBMC and chronic phase in HLN, and therefore the DC-NK cross-talk pathway findings in both studies could be an indication of disease progression from activation of DCs in the acute phase, potentially caused by recognition of parasitic RNAs by TLR9, followed by an inhibition of DC activity in the chronic phase. The involvement of DCs and NK cells in response to *F. hepatica* is a novel finding in both our study and the study on response to *F. hepatica* in HLN (21). DCs and NK cells are present in the liver as a part of the resident immune cell population (71). Therefore, the involvement of these cell types in the response of PBMC and lymph node lymphocytes to *F. hepatica* could indicate that these are important cell types responding to the parasite in the liver, and that these cells could be targeted for stimulation of an adaptive immune response.

The aforementioned ovine HLN study used an infected *vs* control analysis of DE genes, and over 5000 DE genes were detected at 16 wpi (21). However, in our PBMC study only 2 DE genes were found in the infected *vs* control analysis at 16 wpi. Therefore, a comparison of the chronic response in PBMC and HLN in the present study and the study by (21) was not possible. In the HLN study, fibrosis was not found in the lymph node tissue of the infected sheep, and the genes *TGFB1* and *COL1A1* typically associated with liver fibrosis found in the previous ovine studies (20; 72) were not DE in the HLN (21). In our PBMC study, these genes were also not DE, even though their expression would be expected to increase due to the “echoing” capacity of PBMC (73). Fibrosis of the liver was not extensive in the sheep from our study, which is confirmed by the lack of significant overexpression of fibrosis-related genes.

### 4.4 Limitations of longitudinal analysis

Larger numbers of DE genes were found in the longitudinal analysis than when the infected and control groups were compared at each time point. However, it is important to note that the *P* values and fold-changes of these longitudinal DE genes relate to a comparison between the infected animals at week 2 or 16 versus week 0. The statistical difference between the infected and control group is not accounted for in this case and rather the change in response over time in each group is assessed, and control animals are accounted for by subtracting DE genes present in control animals from the infected animal analysis. This analysis was performed to highlight the potential of developmental or seasonal changes in both groups of sheep, which may lead to a small number of DE genes when infected and control animals are compared. It also highlights the necessity of a suitable control in transcriptomics experiments, where changes in gene expression may be caused by factors other than the infection studied.

The infected *vs* control analysis identified genes where expression difference were high enough enough to overcome individual animal differences between the infected and control animal groups. The low number of DE genes found in the main infected *vs* control analysis also indicates that the systemic response to *F. hepatica* was muted, since larger numbers of DE genes were found in the lymph nodes of the same animals at 16 wpi (21).

### 4.5 Overall impact of the study

The development of new strategies for the control of fasciolosis in ruminants is urgent, given the now widespread resistance among liver fluke populations to the only drug effective in killing early immature flukes, and hence protecting against acute fasciolosis in sheep (74). Vaccination is perhaps the most widely-researched and powerful strategy. However, despite decades of focussed research by a number of research groups, significant commercial interest, and limited success in the use of recombinant excretory-secretory proteins from the parasite as vaccine antigens, reviewed by (14) there is as yet no consistent evidence to support the level of vaccine-mediated protection required to bring a vaccine to market. Given the complexity of the target, its complicated life-cycle, lack of immunity from natural infection (56) and propensity to subvert host immune systems (55), these difficulties are in themselves not surprising. In order to overcome the considerable hurdles involved, it may be necessary to adopt novel approaches. Given the difference in disease phenotype between cattle (chronic disease only, economic losses but only very occasional acute disease or mortality) and sheep (acute disease with high morbidity and mortality not unusual, effectively now limiting areas that can be grazed by sheep)(11), one such novel approach might be to ask why the nature of disease varies so much between these two hosts, and whether a “smart vaccine” strategy could, if falling short in terms of rendering sheep resistant to re-infection, could at least limit the pathology to the range of that seen in cattle, thus eliminating mortality and acute morbidity. Exploring the differences in the response of these two species in the acute stage of infection, as we have done in this transcriptomic study, has the potential to indicate strategies, such as use or avoidance of particular adjuvants, and/or co-administration of immunomodulating compounds, that might be useful in this respect.

### 4.6 Concluding remarks

This is the first study which describes the transcriptomic response of ovine PBMC to *F. hepatica* with infection duration up to 16 wpi and with control animals included in the experimental design. The study was systematically compared to a similar study in cattle, both of which represent major livestock species that are the targets for development of vaccines as a control measure for fasciolosis. Overall, the infection *vs* control analysis in our study revealed downregulation of apoptosis and increased cellular metabolism in the acute phase, with the upregulation of the TP53 pathway, and a response to intracellular foreign nucleic acids through TLR9 and RIG-I in the acute phase of infection. The intracellular immune response to RNA molecules could indicate involvement of *F. hepatica* extracellular vesicles in triggering the acute phase response. Based on the most significant DE genes including *CD83*, expression of *TLR9* and a significant DC-NK cross-talk pathway, the involvement of DCs may be key to the acute phase response in sheep. A further longitudinal analysis revealed that an anti-inflammatory response tends to occur in sheep only in the chronic phase of infection, while in cattle the same response is activated as early as 1 wpi. This reflects the different phenotype of disease in the two host species, and may point to new control strategies for sheep, such as a potential benefit of using adjuvants in ovine vaccines to trigger an early anti-inflammatory response, or triggering dendritic cell activation. Overall, this study leads to better understanding of host-parasite relationships.

Breed differences and the infecting strain may have affected the DE genes found in this study when compared to other studies. Furthermore, it is apparent that differences in experimental design and bioinformatic approaches used, including selection of controls, use of software packages, or inclusion of batch effects can significantly affect the resulting DE genes. It is recommended that uninfected control animals are used in the experimental design to account for potential seasonal or developmental differences, especially in long term studies. Use of different sheep breeds as well as different *F. hepatica* strains would be desirable in follow-up studies to validate the findings of our study.

## 5 Data Availability Statement

The datasets generated for this study can be obtained from the European Nucleotide Archive (www.ebi.ac.uk/ena) with accession PRJEB45790.

## 6 Ethics statement

This experiment was approved by the Bioethics Committee of the University of Córdoba (UCO, Spain) (code No. 1118) and conducted in accordance with European (2010/63/UE) and Spanish (RD 1201/2005) Directives on animal experimentation. Written informed consent was obtained from the owners for the participation of their animals in this study.

## 8 Author Contributions

DAN analysed the data and drafted the manuscript. AN-L performed laboratory work. VM-H, AM-M and JP conceived and designed the experimental trial. VM-H collected samples from animals and carried out PBMC isolation. JB supported laboratory work. DEM supported bioinformatics analysis. GM contributed to the writing of the manuscript. DAN, AN-L, VM-H, JP, GM, and DEM contributed to the editing of the manuscript critically for important intellectual content. All authors read and approved the final manuscript.

## 10 Acknowledgments

We would like to thank the staff of the experimental facility from the Universidad de Córdoba for the care of the animals and help provided during the trial.

This article previously appeared as a pre-print on biorXiv, doi: https://doi.org/10.1101/2021.06.21.449266

