## Supplementary Figures for "Timing of transcriptomic peripheral blood mononuclear cell responses of sheep to *Fasciola hepatica* infection differs from those of cattle, reflecting different disease phenotypes"

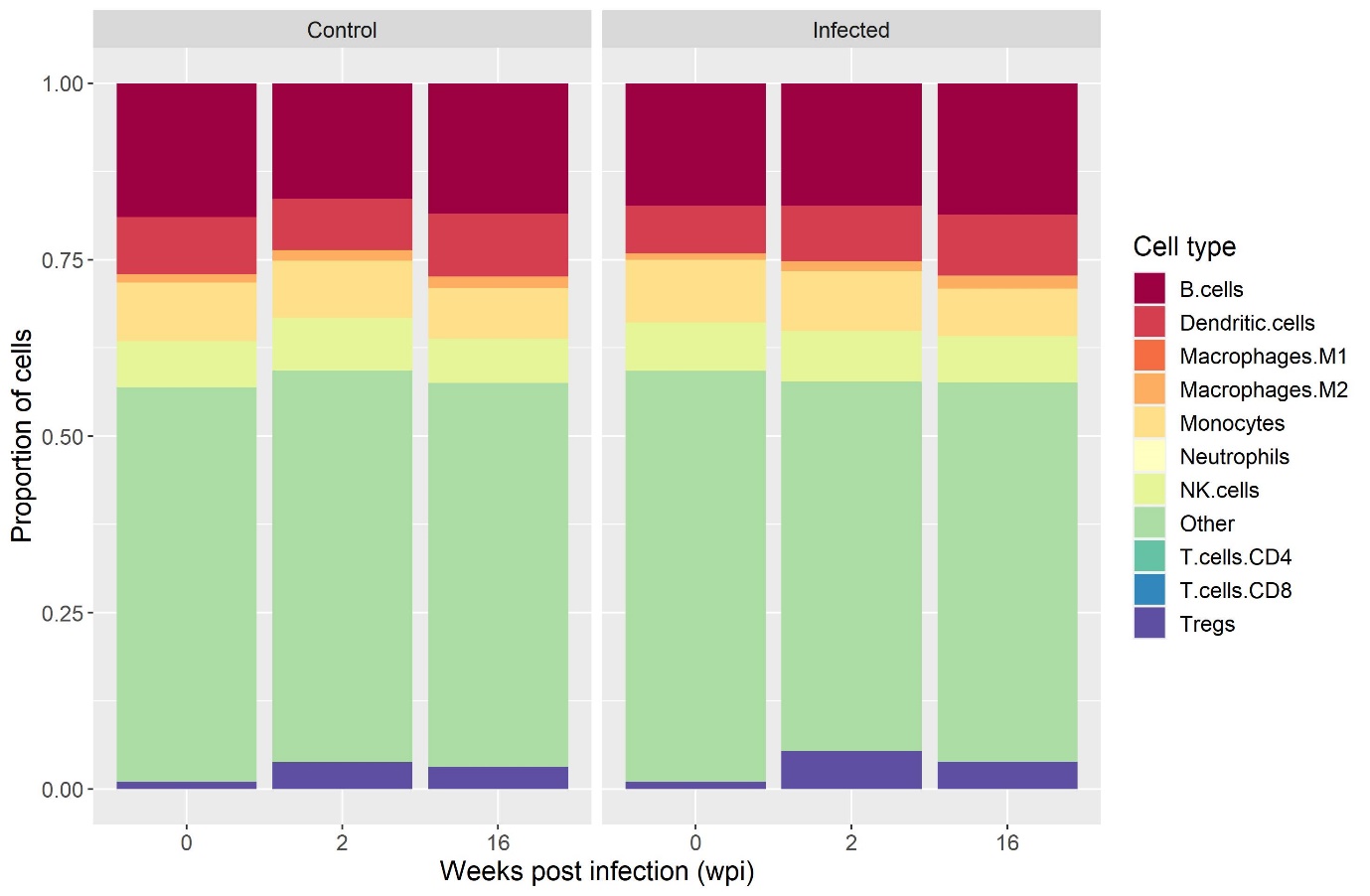

**Supplementary Figure 1.** Changes in cellular composition in PBMC infected and control groups over time illustrated by cell proportions.

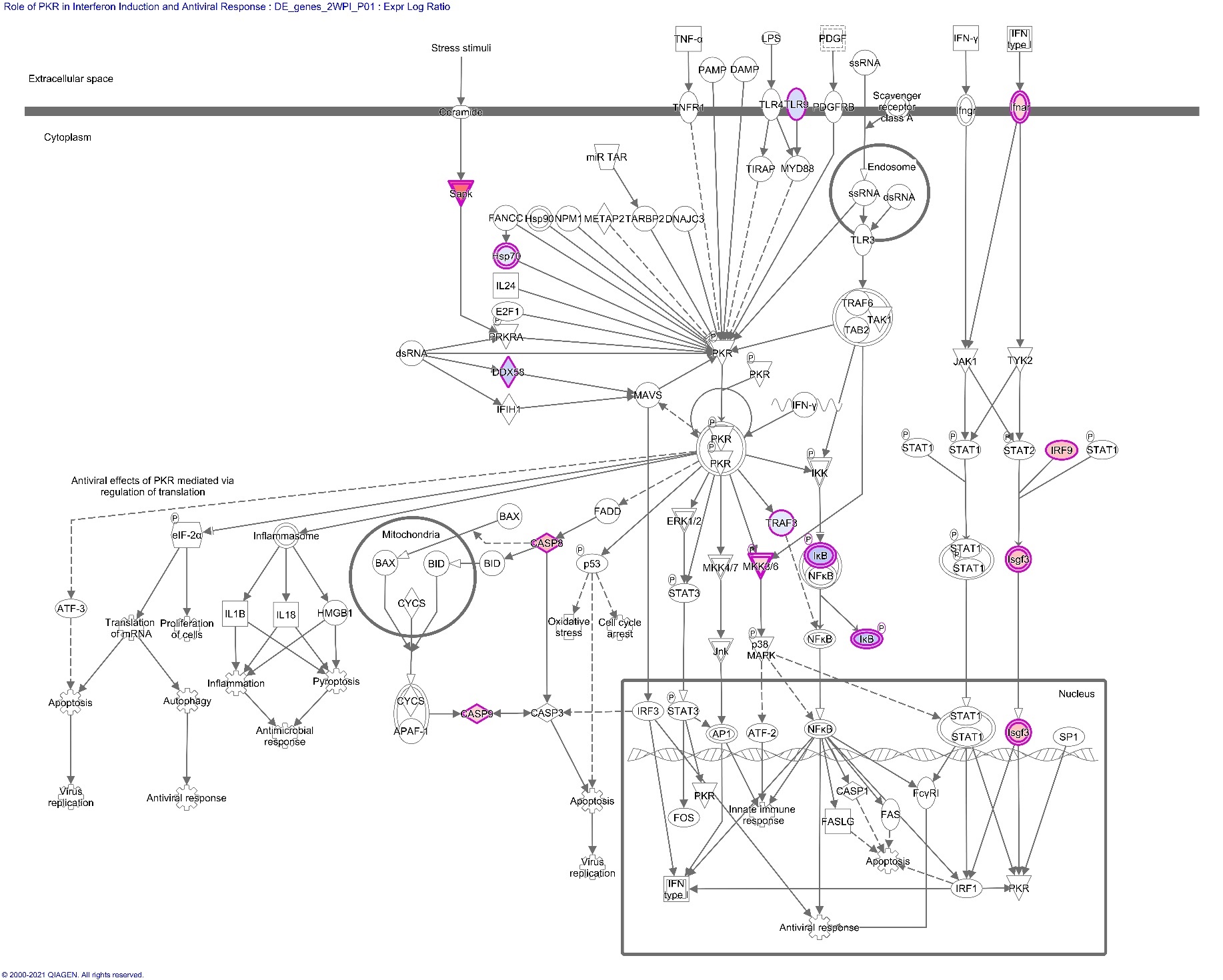

**Supplementary Figure 2**. Role of PKR in interferon induction and antiviral response pathway and the genes expressed in PBMC involved in the pathway at 2 wpi. Upregulated genes are coloured in blue, while downregulated genes are coloured in red. Transcription factors and key genes are denoted with a double-lined outline. This is an inhibited pathway which is involved in response to viral particles in the intracellular space and lead to production to interferons. Among the DE genes involved in this pathway, antiviral receptors and the transcription factor TRAF3 are up-regulated, while the downstream genes involved in cellular apoptosis (caspases) and interferon production (ifnar and IRF9) are downregulated.

Gene names for DE genes involved in this pathway: *CASP8* - caspase 8, *CASP9* - caspase 9, *DDX58* – RIG-I receptor, Hsp70 (*HSPA5)* - heat shock protein family A (Hsp70) member 5, ifnar (*IFNAR2)* - interferon alpha and beta receptor subunit 2, IκB (*NFKBID)* - NFKB inhibitor delta, *IRF9* - interferon regulatory factor 9, Isgf3 – ISGF3 transcription factor, formed by a complex between IRF9 and STAT1:STAT2, MKK3/6 (*MAP2K3)* - mitogen-activated protein kinase kinase 3, Sapk (*MAPK11)* - mitogen-activated protein kinase 11, *TLR9* – Toll-like receptor 9, *TRAF3* - TNF receptor associated factor 3.

**Supplementary Table S1**. Sample metadata and parameters of sequencing and alignment quality.

**Supplementary Table S2.** Log_2_ fold changes and adjusted *P* values of all differentially expressed genes (Benjamini‐Hochberg adjusted *P* value < 0.1) in response to *F. hepatica* at 2 wpi - infected vs control analysis. NA - gene not differentially expressed.

**Supplementary Table S3**. All significantly overrepresented gene ontology, KEGG and REACTOME pathways in response to *F. hepatica* at 2 wpi. The genes were converted to human orthologs and sorted by adjusted P value.

**Supplementary Table S4.** IPA canonical pathways in PBMC in response to *F. hepatica* infection in sheep at 2 wpi – Infected vs Control DE gene analysis.

–log*P* - The negative log_10_ of the probability of association of genes from our data set with the canonical pathway by random chance alone using Fisher’s exact test.

ratio - The number of genes in our dataset that meets the cut-off criteria (P*_adj._* < 0.1), divided by the total number of genes involved in that pathway.

z-score - Overall activity status of the pathway. Z < 0 indicates a prediction of an overall decrease in activity while Z > 0 predicts an overall increase in the activity. NA indicates pathways that are currently ineligible for a prediction.

**Supplementary Table S5.** Log_2_ fold changes and adjusted P values of all differentially expressed genes (Benjamini‐Hochberg adjusted *P* value < 0.05) in response to *F. hepatica* at 2 and 16 wpi - longitudinal analysis. NA - gene not differentially expressed.

**Supplementary Table S6.** Common DE genes in PBMC at the chronic stage of *F. hepatica* infection in sheep and cattle.

**Supplementary Table S7.** IPA canonical pathways in PBMC at acute and chronic stages of *F. hepatica* infection in sheep and cattle.

–log*P* - The negative log_10_ of the probability of association of genes from our dataset with the canonical pathway by random chance alone using Fisher’s exact test.

z-score - Overall activity status of the pathway. Z < 0 indicates a prediction of an overall decrease in activity while Z > 0 predicts an overall increase in the activity. NA indicates pathways that are currently ineligible for a prediction.
